# Sensory and cognitive factors affecting multi-digit touch: a perceptual and modeling study

**DOI:** 10.1101/2021.02.25.432852

**Authors:** Irena Arslanova, Shinya Takamuku, Hiroaki Gomi, Patrick Haggard

## Abstract

Whilst everyday interactions with objects often involve multiple tactile contacts, integration of tactile signals remains poorly understood. Here we characterise the integration process of tactile motion on multiple fingerpads. Across four experiments, participants *averaged* the direction of two simultaneous tactile motion trajectories delivered to different fingerpads. Averaging performance differed between within- and between-hands in terms of sensitivity and precision but was unaffected by somatotopic proximity between stimulated fingers. The sensitivity to the average direction was influenced by the discrepancy between individual motion signals, but only for within-hand conditions. This was explained by a model, in which the ‘virtually leading finger’ received a higher perceptual weighting. Precision was greater in between-hand compared to within-hand conditions. While biased weighting accounted for differences in sensitivity, it was not sufficient to explain the difference in precision, implying additional sensory limitations during within-hand integration. We suggest that unimanual integration is limited and thus exploits a ‘natural’ cognitive prior involving a single object moving relative to the hand to maximise information gain.

**Author summary:** Tactile stimulation is always on. Yet little is known about how the brain combines widespread tactile inputs for perception. Most tactile studies emphasize a single point of tactile stimulation (e.g., location or intensity of a static stimulus) and minimal units of tactile perception (e.g., acuity or selectivity). However, our daily interactions with the world involve encoding spatially and temporally extended tactile signals. Perceiving tactile objects and events as coherent entities requires the somatosensory system to aggregate tactile afferent signals across separate skin regions (i.e., separate digits). Across four experiments we asked participants to *average* direction of two tactile motion trajectories delivered simultaneously to two different fingerpads, either on the same, or on different hands. Our results show strong integration between multiple tactile inputs, but subject to limitations for inputs delivered within a hand. Our model suggests that tactile inputs are weighted according to an integrative model of hand-object interaction that operates within-hands on purely geometric information to prioritise ‘novel’ information from a ‘virtually leading finger’ (VLF).

## Introduction

When an object moves across the skin, as when manipulating an object with the fingers, each finger receives distinct information about object movement. A unified experience of the object’s motion arises due to the brain's ability to integrate individual afferent inputs across spatially discontinuous parts of the receptor surface (the skin). While several studies have revealed mechanisms of tactile motion perception (Pei et al., 2008; 2010; 2011; Pei and Bensmaia, 2014; Pack and Bensmaia, 2015), these investigated single digit stimulation. Thus, multi-digit tactile motion integration remains poorly understood. In a previous study (Arslanova et al., 2021), we found evidence for a neural mechanism that promotes multi-digit tactile integration by modulating the degree of suppressive interactions between stimulated digit representations. Here, we sought to characterise two aspects of this integration process – sensory-integration performance and sensory weighting among digits – and examined whether the latter aspect was based on natural statistics of tactile motion stimuli.

Multi-digit tactile integration is not a straightforward task. Tactile information processing appears limited to two or three simultaneous stimuli (Driver and Grossenbacher, 1996; Gallace, Tan, & Spence, 2006; for review see Spence and Gallace, 2007). Indeed, previous studies have shown that participants are unable to reliably estimate the total intensity or frequency of tactile stimuli spanning separate skin regions (Ho et al., 2011; Walsh et al., 2016; Cataldo et al., 2019; Kuroki et al., 2017). Interestingly, such capacity limitations seem to be more severe when stimuli are delivered to two fingers of the same hand, compared to different hands (Craig, 1985). Previous studies (Walsh et al., 2016; Cataldo et al., 2019) have also revealed that stimuli delivered to multiple fingers of the same hand are often integrated with unequal sensory weights, but not when they are delivered to two fingers on opposite hands. Capacity limitation in general enforces us to allocate available resources to more relevant stimuli. Such resource allocation can be based not only on the reliabilities of individual cues (Ernst and Banks 2002; Oruc et al., 2003; Ernst and Bülthoff, 2004; Alais and Burr 2004) but also on the novelty of given information (Yang et al., 2018) to maximise information gain (Friston et al. 2015; Itti and Baldi 2009; Vergassola et al. 2007). These strategies can lead to biases in sensory weighting.

Natural statistics of tactile motion stimuli suggests a particular strategy for sensory weighting. Different fingers of the *same hand* often receive redundant information. For instance, when an object slides across multiple fingers, one of the fingers will receive a novel stimulus event, such as an edge of the object or a surface texture change, first (we call this the ‘*virtually leading finger*’). The remaining finger(s) would then receive similar stimulation, but with a slight temporal delay. If the brain prioritises *novel* information, the virtual leading finger will attract more attention and have a greater impact on the overall perception, but only when multi-digit stimuli are integrated. We term this *virtual-leading-finger priority (VLF-priority)*.

A VLF-priority explanation is consistent with previous literature on leading fingers in humans and leading whiskers in rats. For example, in Ziat et al.’s (2010) study participants ran their index and middle fingers across a Braille dot stimulator. When the dot location was unpredictably changed after the first finger touched that dot, participants failed to detect the change. This suggests that tactile information was suppressed on the following finger after the leading finger had already touched the stimulus. Drew and Feldman (2007) showed that when rat whiskers were stimulated in a sequence, the neuronal response was strongest for the stimulation on the leading whisker. However, in both these examples, the finger (or whisker) that was virtually leading and thus prioritised depended on *temporal priority*.

Nevertheless, temporal sequence of finger stimulation in natural interactions is tightly coupled with the *geometric* relationship between the fingers and the relative motion of the touched object. Therefore, even when fingers are stimulated synchronously, without any phase difference between stimulations, a virtual leading finger can still be defined by considering the direction of motion of an implied virtual object that causes the stimulations. The VLF-priority hypothesis predicts that the brain will automatically prioritise one finger based on the purely *geometric* relationship between the fingers and the tactile motions, even when tactile stimuli are not in fact caused by a single coherent object motion. Prioritisation of a virtual leading finger under such somewhat artificial conditions provides a powerful laboratory demonstration of how the brain uses an implicit prior of hand-object interaction in perceiving tactile motions.

Here, we characterised the ability to integrate independent tactile motion signals into a percept of overall motion direction based on four experiments. We delivered two tactile motion stimuli (point stimulus movement across skin) simultaneously to two fingerpads, and asked participants to estimate the *average* direction of those two trajectories. Using two separate stimuli allowed us to manipulate the discrepancy between the directions of the individual trajectories to examine how averaging ability is constrained by the distribution of component inputs. We also varied the digits to which the tactile stimuli were presented. In experiments 1 and 3, the stimuli were delivered to fingers on the same hand, whereas in experiment 2 and 4, the stimulated fingers were on different hands. In addition, in experiment 3 and 4 we manipulated the somatotopic relationship between stimulated fingers (Fig. 1A). Although the hand itself was static, the combined tactile motion signals were consistent with implied movement of an object relative to the hand in a different direction on each trial, and consequently defined the ‘virtual leading finger’. Because the direction of tactile motions delivered to each finger was systematically varied across trials, it was possible to infer the relative contribution (i.e., weight) of each finger in making the average direction judgement. This enabled us to test the VLF-priority hypothesis using the collected data. Model analyses further examined whether the VLF-priority hypothesis explained the behavioral patterns observed in the experiments.

**Figure 1.**
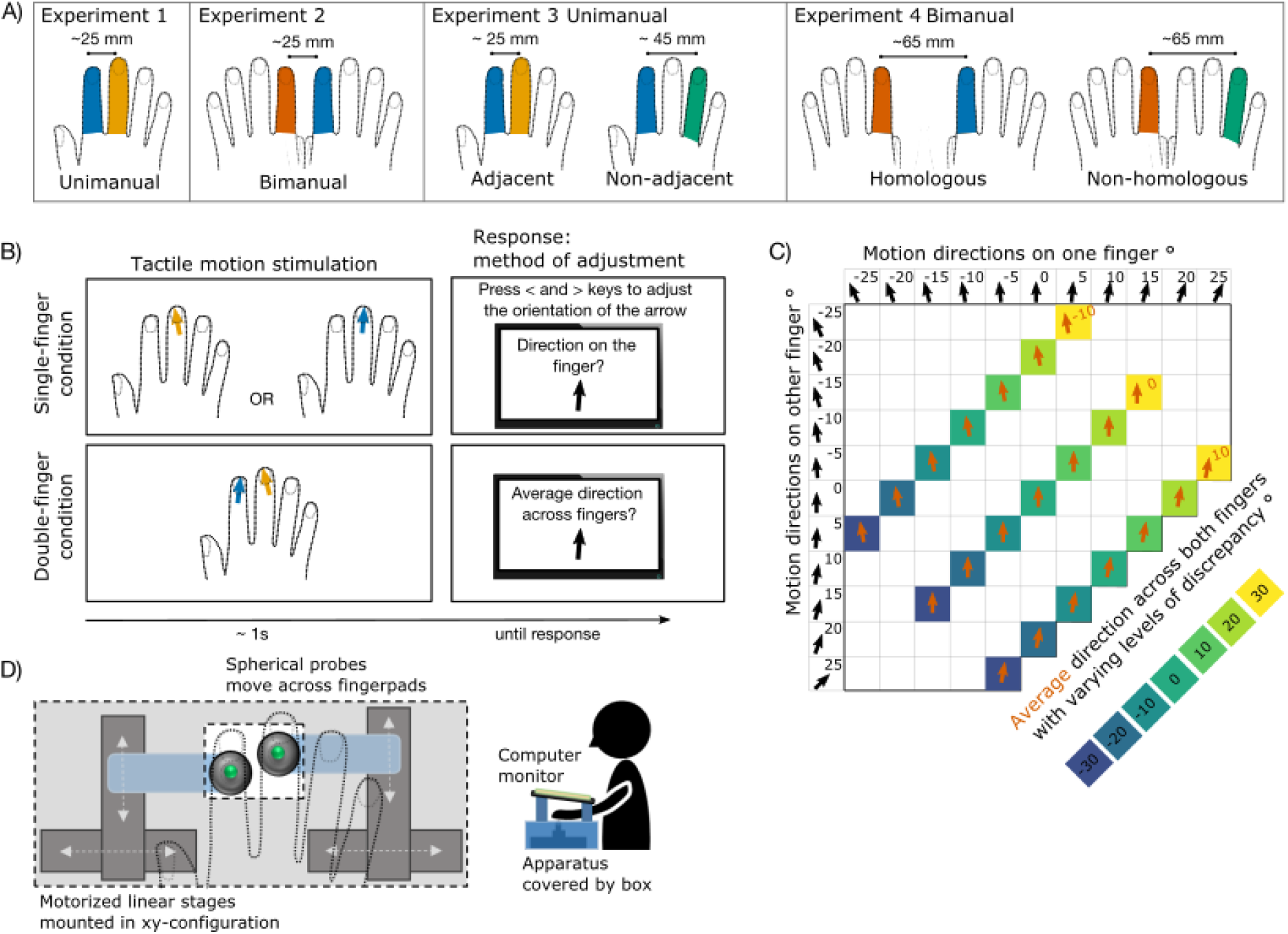
Methods. **A)** Stimulated fingers in each experiment and condition. **B)** Paradigm and trial example from Experiment 1. Fingers were stimulated in two main conditions: single-finger condition and double-finger condition. In single-finger condition, in different blocks, just the index or just the middle finger was stimulated. Participants reported the direction of the stimulus’ movement across their fingerpad. To do so, they adjusted a visual pointer on the screen after each trial (by pressing left and right keys or left and right foot pedals). In double-finger condition, both component fingers were stimulated simultaneously, and participants had to judge the average direction between the two component directions. **C)** Tactile stimuli. Eleven component single-finger directions were combined into 3 average directions with 7 levels of discrepancy. The sign of discrepancy reflects whether the component directions tended to converge (negative discrepancy) towards the inner edges of fingertips or diverge (positive discrepancy) towards the outer edges of fingertips. **D)** Tactile motion apparatus and set-up. Tactile apparatus consisted of two spherical probes (4 mm diameter) attached to two motorized linear stages mounted in a XY-axis configuration. The apparatus was covered by a wooden box with a rectangular gap, which was used to guide finger placement and secure the hand position. A computer screen, where participants indicated their response, was placed above the hand.

## Results

To investigate the aggregation ability, we compared perception of the average direction (double-finger condition) to perception of the two component stimuli presented alone (single-finger condition, see Fig. 1B). In single-finger conditions, only one probe was moved, touching only one fingertip, and participants had to estimate the direction of its movement. In double-finger conditions, both probes were moved simultaneously along both fingertips, and participants had to estimate the average direction of the two trajectories. We used 21 possible combinations (Fig. 1C) that produced three different average motion patterns, with varying discrepancy between the two motion directions. The sign of discrepancy reflects whether the component directions tended to converge towards the inner edges of fingertips (negative discrepancy) or diverge towards the outer edges of fingertips (positive discrepancy). Participants gave their response after the motion stimuli, by adjusting the orientation of a visual arrow that appeared on a monitor placed above their fingertips (Fig. 1D). The conditions were blocked and counter-balanced. Each experiment contained a separate group of 15 participants.

Tactile motion processing was characterised in terms of sensitivity to changes in stimulus direction, bias of direction judgments, and precision of repeated estimates of direction for the same stimulus (Fig. 2; see Methods). Figure 2B shows the fitted linear regression to a representative participant in Experiment 1. The slope of the regression reflects the relationship between perceived direction and actual tactile direction, while the intercept reflects the perceptual shift of the perceived midline of a given finger. We expected the slope to be greater than 0 and close to 1, reflecting participants’ ability to perceive spatiotemporal information from the skin. Because we did not manipulate finger posture, and always aligned the finger long axis with the 0° stimulus direction, we expected intercept values to be close to 0 reflecting unbiased perception. A perceiver with a given level of sensitivity or bias may be more or less precise (Fig. 2A). We used unbiased standard deviation (SD; see Methods) of repeated judgments of the same stimuli as a measure of precision.

**Figure 2.**
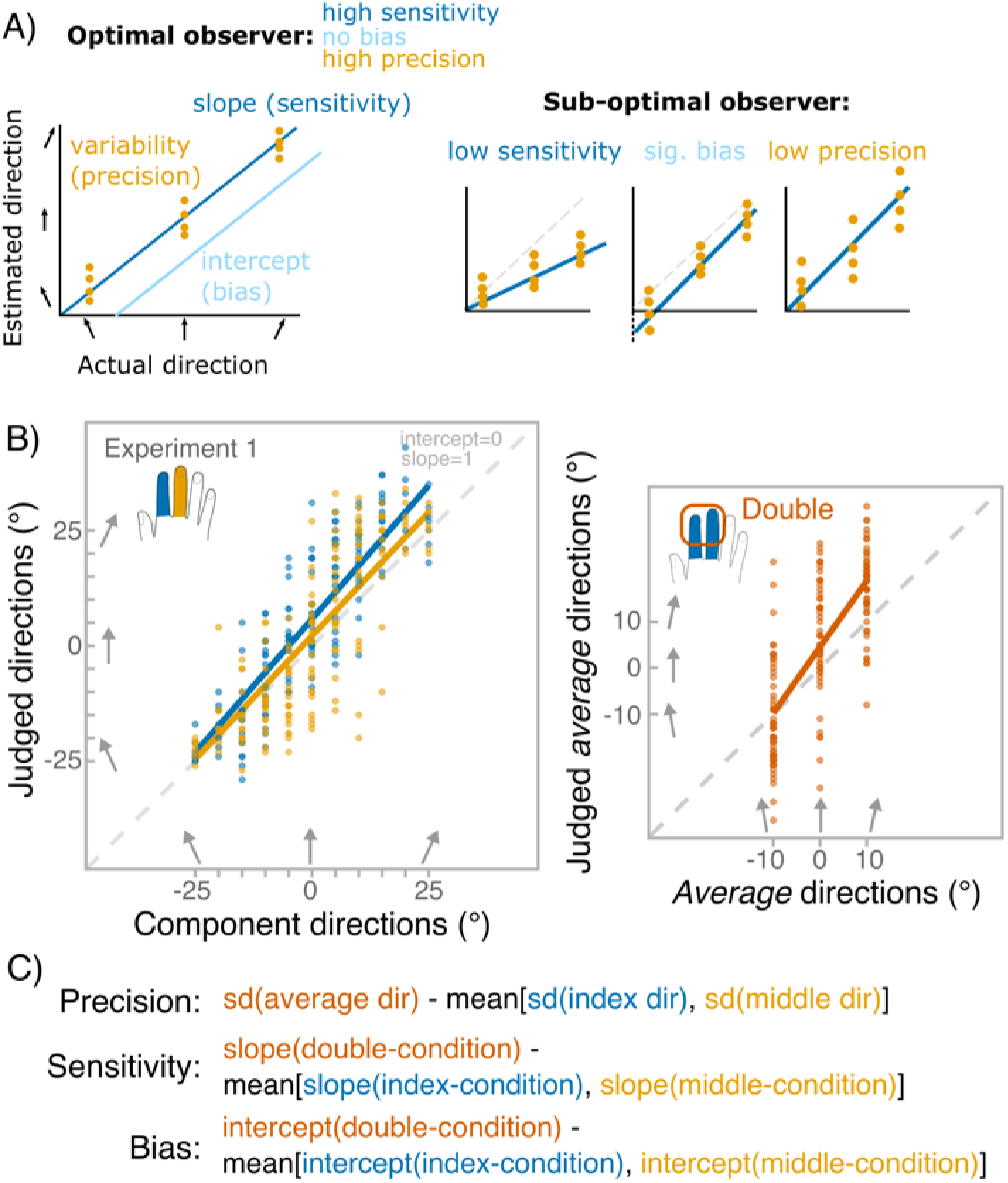
Analysis. **A)** Measures used to characterise tactile motion perception: sensitivity, bias and precision. Sensitivity quantifies the ability to perceive differences between motion directions. Bias reflects the shift of the perceived 0° motion, in our case corresponding to the midline of the finger. Precision (also termed reliability) measures the consistency of a direction percept elicited by repetitions of the same stimuli. The schematic panels show that these three measures are independent. **B)** Data from an example participant in Experiment 1. Linear regression was fit to the data separately for the two single-finger conditions (right panel) and the double-finger condition (left panel). Slope and intercept values reflecting sensitivity and bias, respectively, were estimated from the regression fit. The dots indicate repeated judgements of each direction. Standard deviations (SDs) were calculated for each direction reflecting the precision. Note that each average direction was composed of one component direction delivered to the index finger and another delivered to the middle finger resulting in 21 combinations: 7 discrepancy levels for a given average direction. In the left panel plot, the discrepancy has been pooled together. **C)** Calculation of main measures. We were interested in the contrast between perceiving the average direction versus estimating individual component directions. For precision we contrasted SD for average direction to the mean SDs for corresponding component directions, while for sensitivity and bias, we contrasted slope and intercept of double-finger condition to the mean slope and intercept value of both single-finger conditions.

We performed preliminary analyses of each single-finger condition, which are reported in supplementary material (S2). Perceiving the average direction requires participants to integrate the two single-finger component directions. We therefore compared performance for perceiving the average direction with the *mean* perceptual performance when perceiving the direction of a single stimulus in the single-finger conditions (Fig. 2C). Better performance in the double finger condition would imply a benefit of integration, relative to the performance in the corresponding single-finger conditions. Figure 3 shows the fitted linear regressions in each experiment and condition that were used to derive slope and intercept values. Table 1 shows the overview of the main measures. In all experiments, participants showed slope values that were significantly higher than 0 and close to 1. Interestingly, in single-finger conditions the slope was often above 1. Except in Exp.1 and in the homologous condition of Exp.4, perception was generally unbiased. For the main analysis, table S1 shows the full breakdown of all statistical effects, here, we only describe the main findings.

**Table 1.**
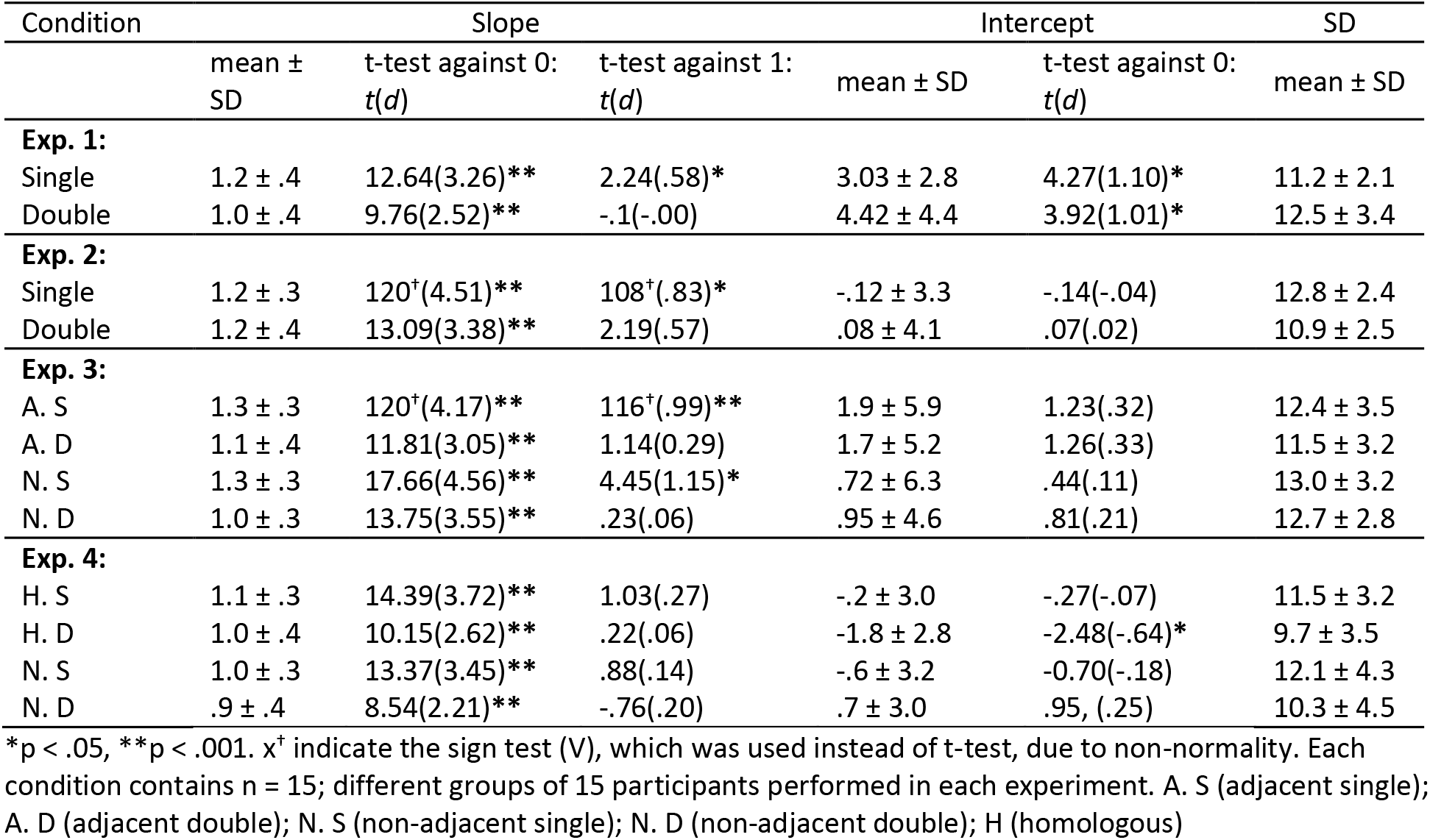
Overview of the main measures across all 4 experiments

**Figure 3.**
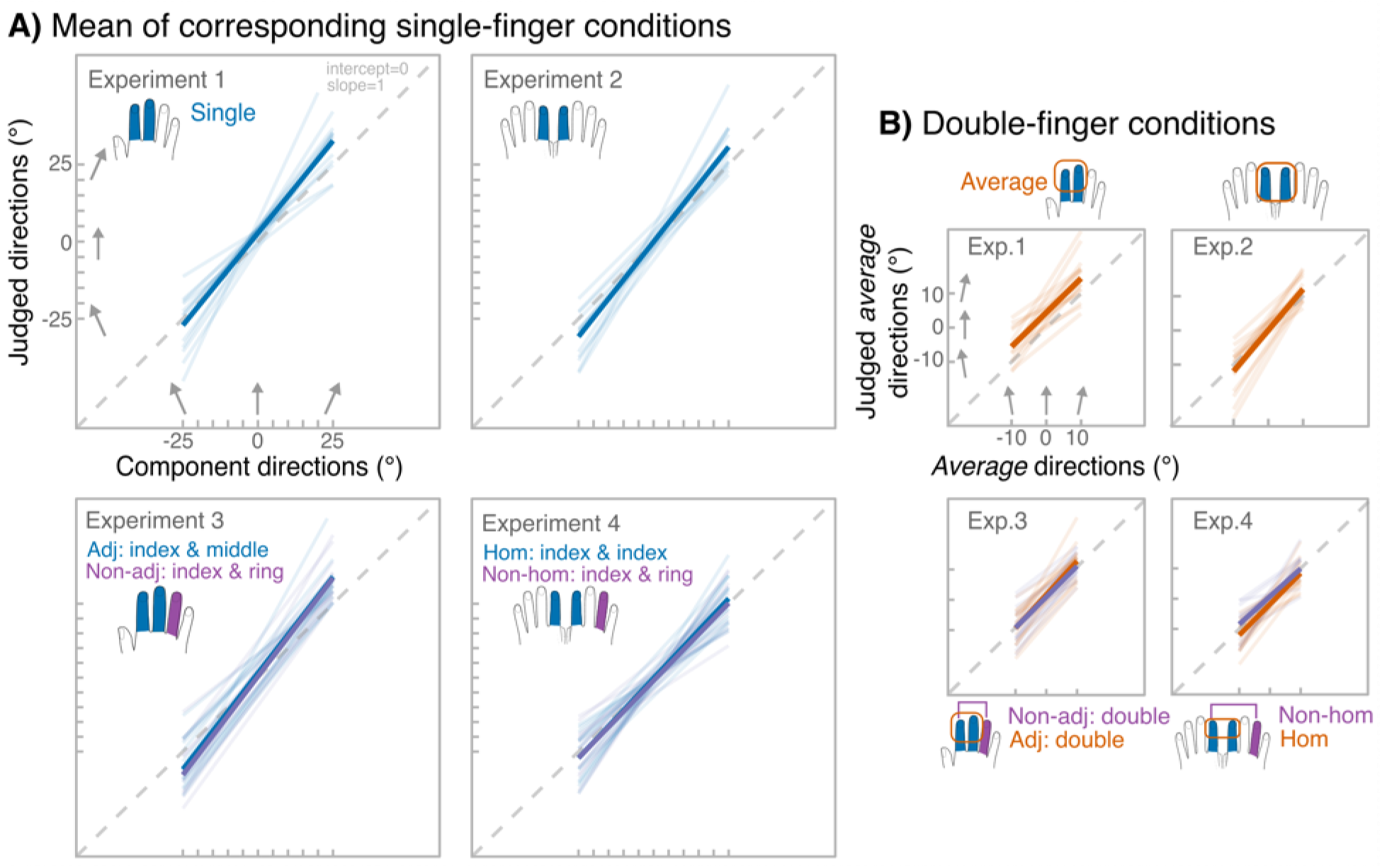
Individual fitted linear regressions (transparent lines) and group-level regressions (thick lines) for each experiment, separately for single-finger conditions (**A**) and double-finger conditions (**B**). The grey dashed line represents the equality line of slope 1 and intercept 0.

### Experiments 1 and 2

Experiment (Exp.1 unimanual vs. Exp.2 bimanual) was a between-subjects factor to test whether averaging ability differed between unimanual and bimanual tasks. Specifically, we focus on the statistical significance of experiment by number-of-finger condition (double vs. single) interaction that would indicate whether averaging ability differed across unimanual and bimanual experiments. We did not find a significant interaction for sensitivity (*F*_1, 28_ = 2.24, *p* = .15, *ηp^2^* = .07, Fig. 4A) or bias (*F*_1, 28_ = .55, *p* = .46, *ηp*^2^ = .02, Fig. 4A). However, the interaction was significant for precision (*F*_1, 28_ = 11.6, *p* = .002, *ηp^2^*= .29, Fig. 4A). Namely, when directions had to be aggregated across hands, the averaging process resulted in a more precise average estimate (mean SD = 10.9, SD = 2.5) compared to the mean estimate of the component directions (mean SD = 12.8 ± 2.4; paired-sample t-test: t_14_ = 4.1, p = .001, d = 1.1). In contrast, when the same directions were delivered to the fingers of the same hand, aggregation did not lead to significant precision benefit (paired-sample t-test: *t*_14_ = −1.59, *p* = .13, *d* = −.41).

**Figure 4.**
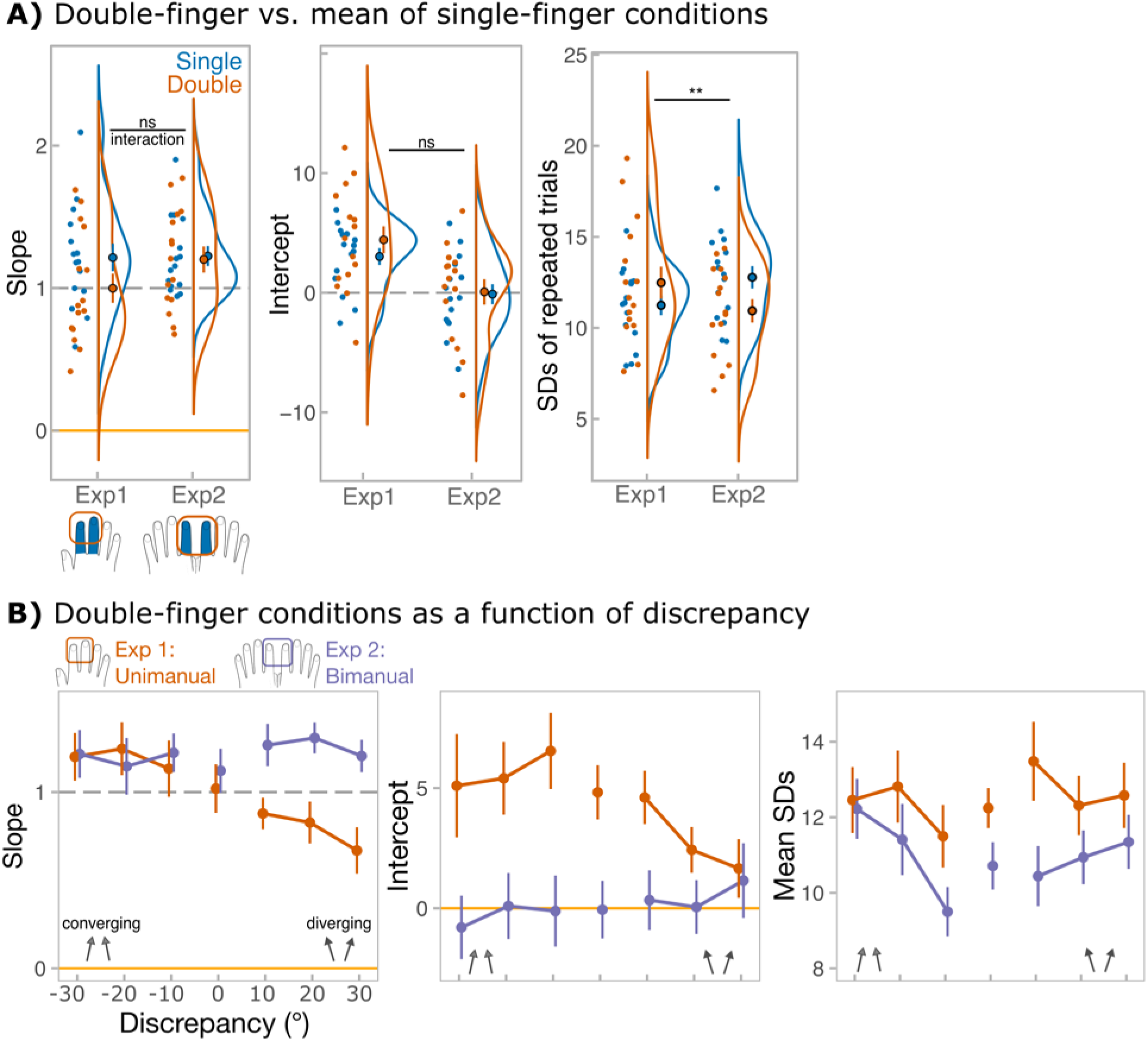
Results of Experiments 1 (unimanual) and 2 (bimanual). **A)** Main measures, from the left: 1) slope values, corresponding to the sensitivity, estimated from single-subject regressions; 2) intercept values, corresponding to the bias, estimated from single-subject regressions; 3) unbiased SD values for repeated estimation of identical directions. SDs were calculated separately for each of 21 stimulus combinations; for simplicity, data was pooled across combinations to show the mean SDs for single-finger (blue) and double-finger (orange) conditions. Points with error bars reflect group-level means and standard errors of the mean. Raincloud plots (Allen et al., 2019) show the distribution of the data. Upper black annotation shows statistical significance of *number of fingers* (single vs. double) by *experiment* (unimanual vs. bimanual) interaction. The data includes two independent groups of 15 participants *per experiment.* **B)** The effect of the discrepancy between the component directions on *averaging* performance. From the left: group-level slope values, group-level intercept values, unbiased SD values for repeated estimations of identical combinations. SDs were calculated for each average direction separately (see Table S1), but for simplicity data was pooled across average directions to show the mean SDs for each discrepancy. In all panels, error bars correspond to standard error of the mean. Discrepancy of 0 (when component directions were identical) is included in the plots for illustrative purposes, but was not included in the analysis, because discrepancy was factored into *sign of discrepancy* (negative discrepancy, when directions converged vs. positive discrepancy, when directions diverged) and *level of discrepancy* (30° vs. 20° vs. 10°). Orange traces reflect unimanual averaging, while purple traces reflect bimanual averaging.

We next explored whether averaging was affected by the discrepancy between component directions. In terms of bias, averaging remained statistically unchanged across both levels of discrepancy and the signs of discrepancy (*p* values were .28 > *p* > .06; Fig. 4B). In terms of precision, there was only a significant interaction between discrepancy level and average direction (*F_4, 112_* = 3.6, *p* = .008, *ηp^2^* = .11). Bonferroni-corrected pairwise tests showed that precision was higher during low discrepancy, but this effect persisted only when average direction was 10°. While sensitivity was not affected by how similar component directions were (main effect of level of discrepancy: F_2, 56_ = .48, p = .62, ηp^2^ = .02; level of discrepancy by experiment interaction: F_2, 56_ = .21, p = .81, ηp^2^ = .007), it was affected by the sign of discrepancy, but only in unimanual experiment (sign of discrepancy by experiment interaction: F_1, 28_ = 11.5, p = .002, ηp^2^ = .29; Fig. 4B). Specifically, during unimanual averaging, the sensitivity dropped when component directions diverged to the outer corners of the fingertips (Bonferroni-corrected pairwise test: *p* < .001; mean slope = 1.19, SD = .56 for converging vs. mean slope = .79, SD = .44 for diverging). No such effect was found during bimanual averaging (Bonferroni-corrected pairwise test: *p* = .45).

### Experiment 3

We examined whether averaging ability changed with somatotopic distance between component directions (Fig. 5). Aggregation in Experiment 1 could have been limited by topographically organised inter-digit interactions, which may result in masking effects (von Békésy, 1967). Because these interactions are believed to follow a spatial gradient (Ishibashi et al., 2000), increasing the somatotopic distance between component directions may affect aggregation performance. For this reason, in Exp. 3 participants had to aggregate directions either across adjacent (index and middle, as in Exp. 1) or non-adjacent fingers (index and ring). Adjacency was a within-participant factor, and because our main interest was whether averaging was modulated by somatotopic distance, we focused on the statistical significance of adjacency by number-of-finger condition interaction. However, none of the measures showed a significant interaction: sensitivity (F_1, 14_ = 1.5, p = .24, ηp^2^ = .10; BF^01^= 1.9 ± 3.3%; BF_incl_ = .5), bias (F_1, 14_ =.16, p = .69, ηp^2^ = .01; BF^01^= 2.8 ± 3.1%; BF_incl_ = .1), and precision (*F*_1, 14_ = .48, *p* = .50, *ηp^2^* = .03; BF^01^= 2.8 ± 4.9%; BF_incl_ = .2). Bayesian factors tended to support the absence of the interaction, showing low evidence for its inclusion into the ANOVA model (BF_incl_ values less than .3 for bias and precision). In terms of discrepancy-dependent effects, as in Experiment 1, sensitivity to average direction dropped when directions started to diverge (main effect of sign of discrepancy: F_1, 14_ = 12.7, p = .004, ηp^2^ = .48; mean slope = .86, SD = .52 for diverging vs. mean slope = 1.26, SD = .58 for converging). This effect was present regardless of the adjacency between the stimuli (sign of discrepancy by adjacency interaction: F_1, 14_ = 1.4, p = .25, ηp^2^ = .09).

**Figure 5.**
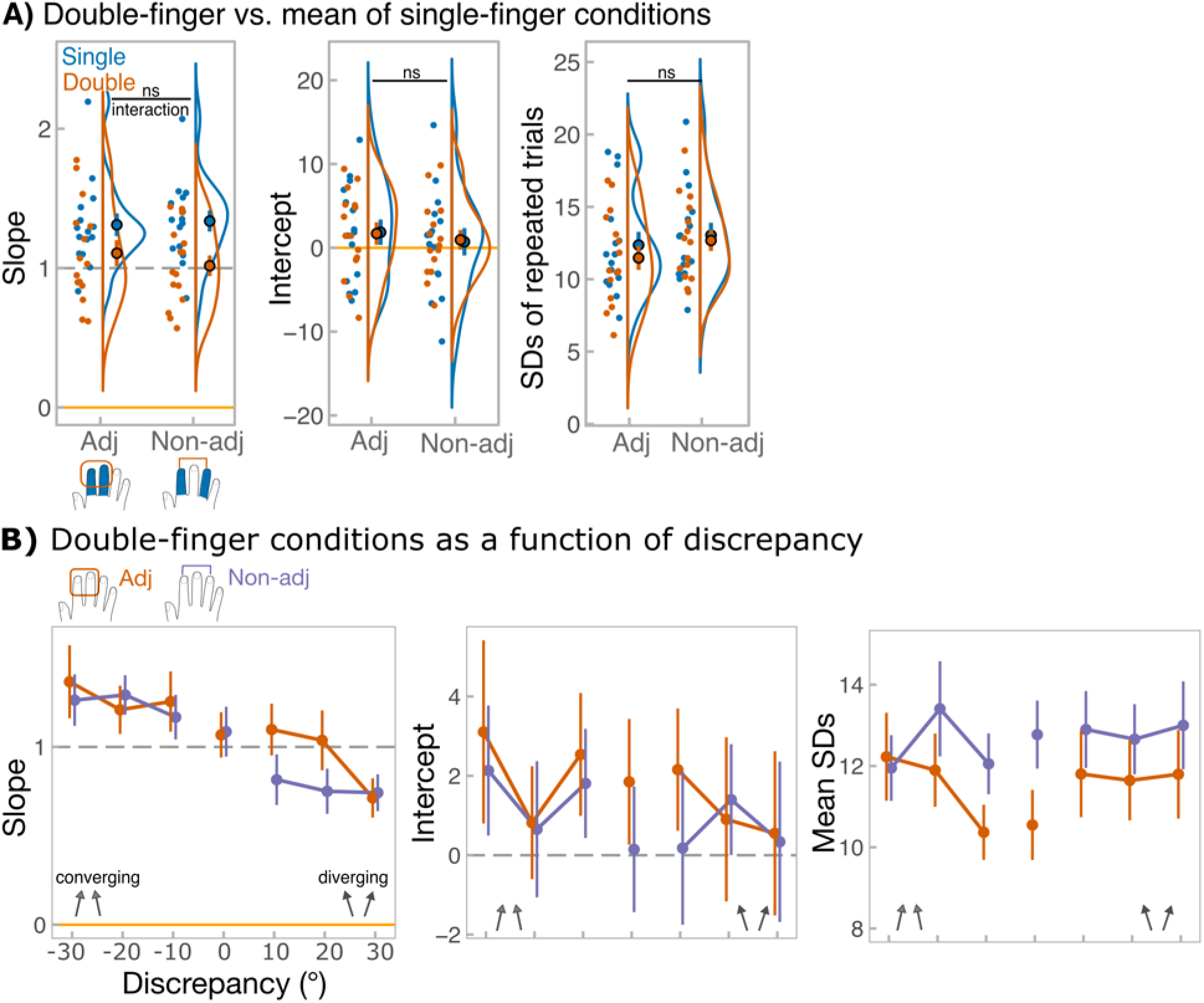
Results of Experiment 3. **A)** Main measures, from the left: 1) slope values, corresponding to the sensitivity, estimated from single-subject regressions; 2) intercept values, corresponding to the bias, estimated from single-subject regressions; 3) unbiased SD values for repeated estimation of identical directions. SDs were calculated separately for each of 21 stimulus combinations, for simplicity, data was pooled across combinations to show the mean SDs for single-finger (blue) and double-finger (orange) conditions. Points with error bars reflect group-level means and standard errors of the mean. Raincloud plots (Allen et al., 2019) show the distribution of the data. Upper black annotation shows statistical significance of *number of fingers* (single vs. double) by *adjacency* (adjacent vs. non-adjacent stimuli) interaction. The data includes the same 15 participants performing under all conditions. **B)** The effect of the discrepancy between the component directions on *averaging* performance. From the left: group-level slope values, group-level intercept values, unbiased SD values for repeated estimations of identical combinations. SDs were calculated for each average direction separately (see Table S1), but for simplicity data was pooled across average directions to show the mean SDs for each discrepancy. In all panels, error bars correspond to standard error of the mean. Discrepancy of 0 (when component directions were identical) is included in the plots for illustrative purposes, but was not included in the analysis, because discrepancy was factored into *sign of discrepancy* (negative discrepancy, when directions converged vs. positive discrepancy, when directions diverged) and *level of discrepancy* (30° vs. 20° vs. 10°). Orange traces reflect averaging over adjacent fingers, while purple traces reflect averaging over non-adjacent fingers.

### Experiment 4

We then examined whether bimanual averaging was dependent on the exact stimulated skin-site. In contrast to strict laterality of primary somatosensory cortex (SI), research has revealed receptive fields in SI that encompass bilateral homologous digits (Iwamura et al. 2002). In addition, tactile judgements from two hands sometimes follow a digit-specific pattern, exhibiting stronger interactions between homologous fingers (for review see Tamè et al., 2016; 2019). Thus, we asked participants to average tactile trajectories either across homologous fingers (as in Experiment 2; left and right index fingers) or non-homologous fingers (left index and right ring; Fig. 6). Again, homology was a within-participant effect and we focused on the statistical significance of homology by number-of-finger condition interaction. As in Exp. 3, no significant interaction was found for sensitivity (F_1, 14_ = .70, p = .12, ηp^2^ = .05; BF^01^= 2.1 ± 9.2%; BF_incl_ = .4) or precision (*F*_1, 14_ = .002, *p* = .96, *ηp^2^* < .001; BF^01^= 2.7 ± 2.8%; BF_incl_ = .4). There was a significant interaction for bias (F_1, 14_ = 6.44, p = .02, ηp^2^ = .32) with significant anticlockwise bias emerging during averaging relative to perceiving component directions individually, but only when stimuli are delivered to homologous fingers (paired-sample simple effect t-test: t_14_ = 2.89, p = .01, d = .75). Again, Bayesian factors tended to support the absence of the interaction. As in Experiment 2, sensitivity to averaged direction was not statistically modulated by how similar component directions were nor the overall direction of the component trajectories relative to the edges of the fingepads (all p values: .38 < p < .96).

**Figure 6.**
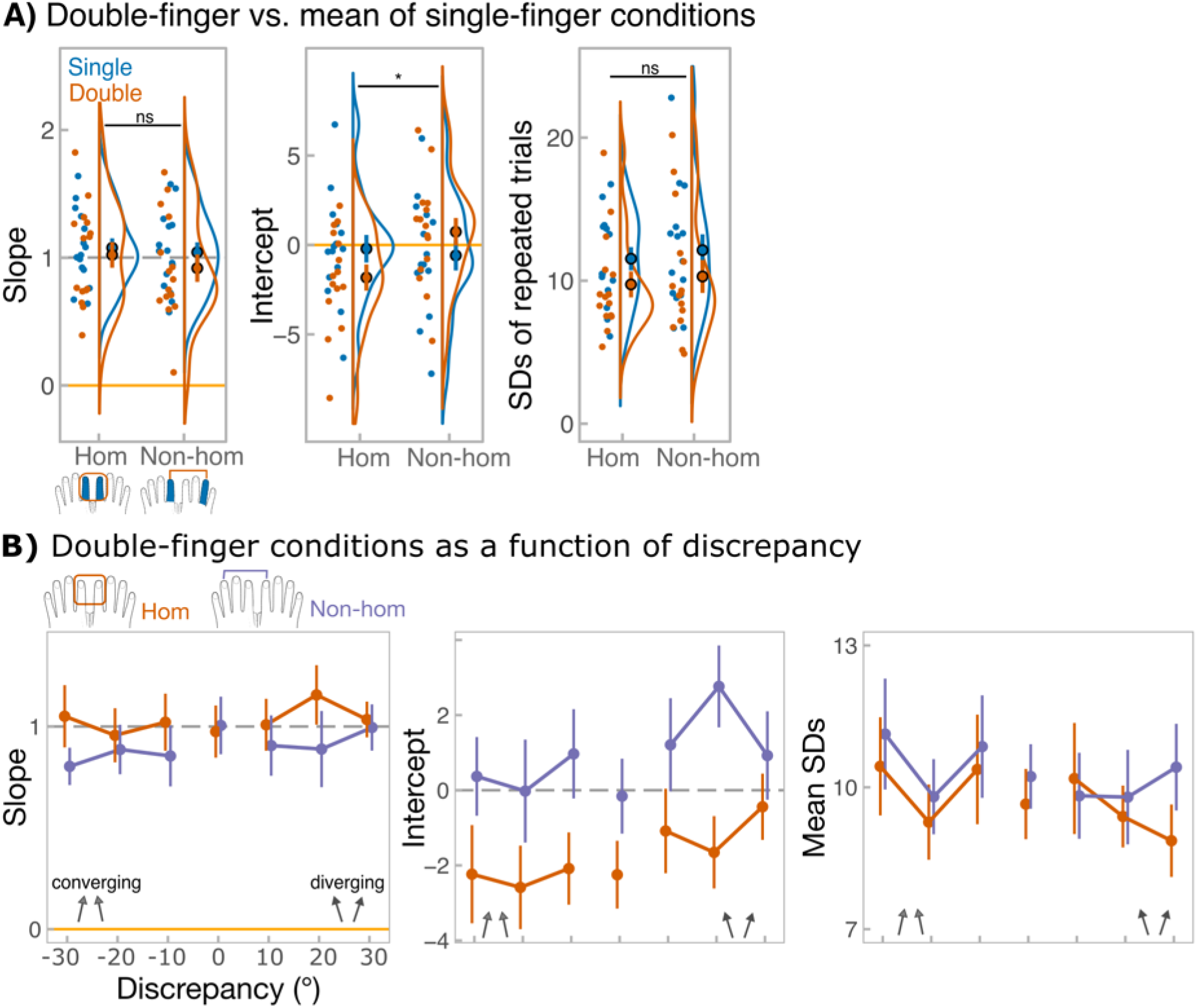
Results of Experiment 4. **A)** Main measures, from the left: 1) slope values, corresponding to the sensitivity, estimated from single-subject regressions; 2) intercept values, corresponding to the bias, estimated from single-subject regressions; 3) unbiased SD values for repeated estimation of identical directions. SDs were calculated separately for each of 21 stimulus combinations, for simplicity, data was pooled across combinations to show the mean SDs for single-finger (blue) and double-finger (orange) conditions. Points with error bars reflect group-level means and standard errors of the mean. Raincloud plots (Allen et al., 2019) show the distribution of the data. Upper black annotation shows statistical significance of *number of fingers* (single vs. double) by *homology* (homologous vs. non-homologous stimuli) interaction. The data includes the same 15 participants performing under all conditions. **B)** The effect of the discrepancy between the component directions on *averaging* performance. From the left: group-level slope values, group-level intercept values, unbiased SD values for repeated estimations of identical. SDs were calculated for each average direction separately (see Table S1), but for simplicity data was pooled across average directions to show the mean SDs for each discrepancy. In all panels, error bars correspond to standard error of the mean. Discrepancy of 0 (when component directions were identical) is included in the plots for illustrative purposes, but was not included in the analysis, because discrepancy was factored into *sign of discrepancy* (negative discrepancy, when directions converged vs. positive discrepancy, when directions diverged) and *level of discrepancy* (30° vs. 20° vs. 10°). Orange traces reflect averaging over homologous fingers, while purple traces reflect averaging over non-homologous fingers.

### Precision across all experiments

To further examine the observed precision benefit from multiple bilateral stimuli over multiple unilateral stimuli, we pooled the data from all four experiments (Fig. 7) and conducted a mixed ANOVA with within-subjects factor number-of-fingers and between-subjects factor experiment. In experiments 3 and 4, we first averaged the responses between adjacency and homology conditions, as these did not significantly differ. The mixed ANOVA yielded a significant interaction between number of fingers and experiment (F_3, 56_ = 4.9, p = .004, ηp^2^ = .21), indicating that the difference in precision between single- and double-finger conditions varied across experiments. As a follow-up analysis, we compared the effect of number of fingers between unimanual (Exp.1 and Exp.3 pooled) and bimanual (Exp.2 and Exp.4 pooled) experiments. It again yielded a significant interaction with number-of-fingers (F_1, 58_ = 10.23, p = .002, ηp^2^ = .15). Specifically, in bimanual experiments, the precision of averaged estimate was enhanced (mean SD= 10.5, SD = 3.2) relative to component direction estimates (mean SD = 12.3, SD = 3.0; main effect of number of fingers when Exp.2/Exp.4 data is pooled: F_1, 28_ = 23.34, p < .001, ηp^2^ = .45; no interaction with experiment: F_1, 28_ < .001, p = .98, ηp^2^ < .001). In contrast, in unimanual experiments, participants did not benefit from averaging (mean SD = 12.28, SD = 3.02) over estimating component directions (mean SD = 12.0, SD = 2.86; main effect of number of fingers when Exp.2/Exp.4 data is pooled: F_1, 28_ = .33, p = .57, ηp^2^ = .01; BF^01^= .9 ± 4.6%; BF_incl=_ .3; no interaction with experiment: F_1,_ 28 = 3.0, p = .09, ηp^2^ = .10). This result supports that averaging produces a more precise or less variable direction estimate, but only when component directions are presented bilaterally.

**Figure 7.**
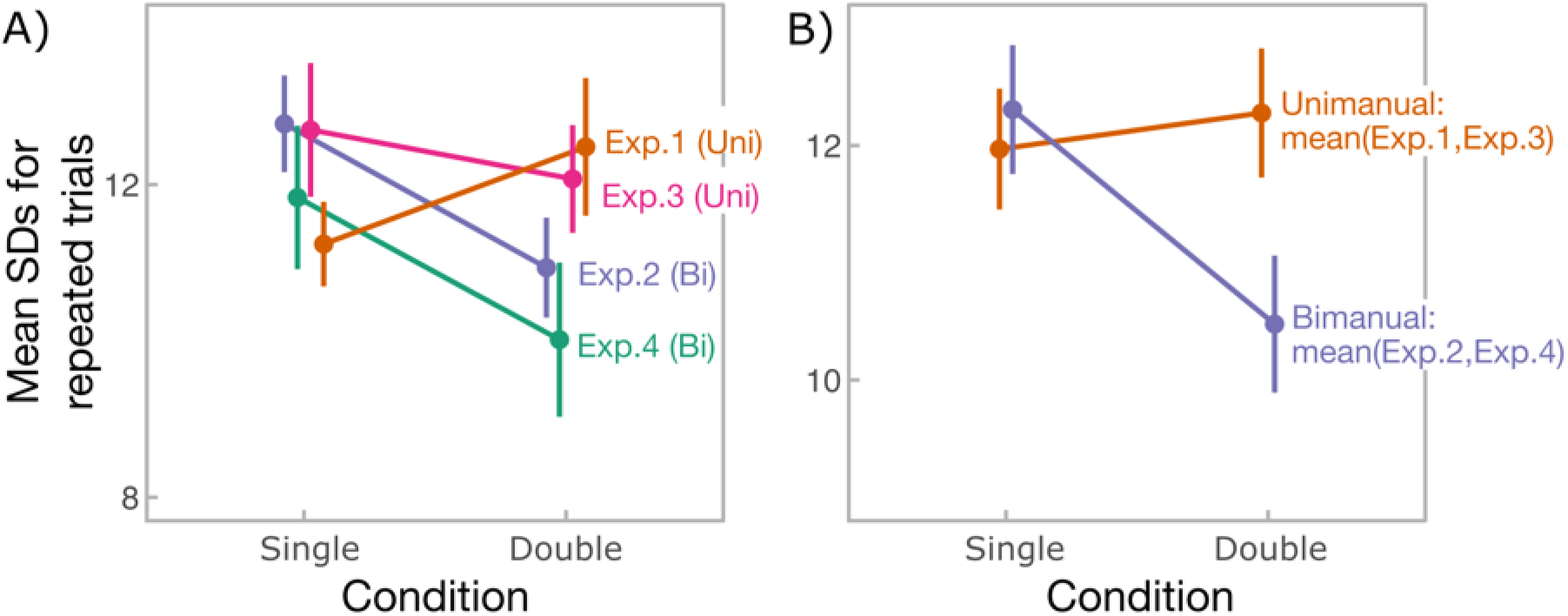
Precision as measured by standard deviations (SDs) for repeated identical trials, separately for every experiment (**A**) and pooled across unimanual and bimanual experiments (**B**). Error bars represent standard error from the mean.

### Sensory weighting model

Behavioral analysis demonstrated distinctive patterns of averaging performance that were consistent across experiments: (1) precision of average direction judgements improved relative to single direction judgements when stimuli were integrated across hands, but not when they were integrated within a single hand; (2) sensitivity to average direction deteriorated as a function of discrepancy between component directions only when stimuli were delivered unimanually. In the following, we tested the VLF-priority hypothesis and examined whether it explained the abovementioned differences between unimanual and bimanual averaging.

We first computed the relative contribution (weighting) of each finger in judging the average direction, directly from the judgements, by assuming accurate estimation of component directions (see Methods for detail). Since the VLF would be defined based on the relative object motion, we expected that the weight depended on the average direction (for breakdown of all effects see Table S3). Fig. 8 shows the computed weight of the left-most finger (right index finger in unimanual experiments and left index finger in bimanual experiments). Indeed, in the unimanual experiments, the weight assigned to the left-most finger changed as a function of average angle both when stimulated fingers were adjacent (F_2, 56_ = 10.0, p = .001, ηp^2^ = .26) and non-adjacent (F_2, 28_ = 6.7, p = .006, ηp^2^ = .32). In contrast, the weight did not depend on the average direction when stimuli were delivered to fingers of different hands, neither when those fingers were homologous (F_2, 56_ = .43, p = .62, ηp^2^ = .02; BF^01^= 35.1 ± .6%; BF_incl=_ .1) nor non-homologous (F_2, 28_ = .33, p = .72, ηp^2^ = .02; BF^01^= 20.4 ± .7%; BF_incl=_ .2). Further analysis of the unimanual conditions showed that the weight was greater for stimuli with average direction of 10° compared to those with average direction of −10° (post-hoc Bonferroni-corrected pairwise tests; adjacent: p= .004; non-adjacent: p= .001; see Fig. 8) in agreement with the hypothesis. The weighting, however, also changed as an interaction between discrepancy level and direction, an effect that was significant for adjacent fingers (F_2, 56_ = 15.7, p = .002, ηp^2^ = .36). This pointed to the potential effect of errors in estimating the component directions (addressed in S4).

**Figure 8.**
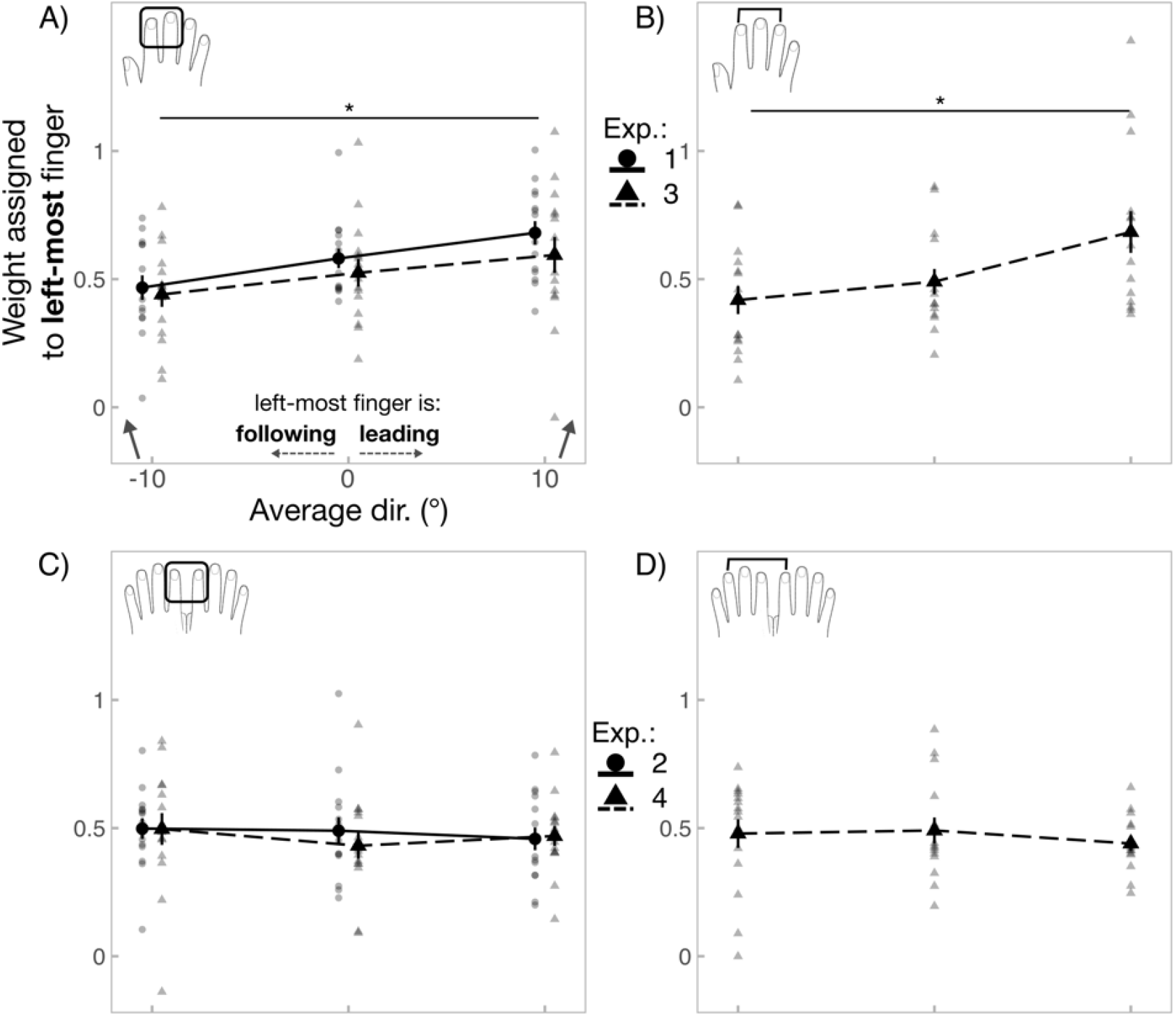
Estimated weight given to the left-most finger in the observed conditions. **A)** and **B)** show results for the unimanual averaging. **A)** compromises pooled data across adjacent conditions (N = 30) and **B)** represents data from the non-adjacent condition (N = 15). **C)** and **D)** shows bimanual averaging results. **C)** contains data from the homologous condition (N = 30) and **D)** represents data from non-homologous condition (N = 15). Left-most finger was the virtual leading finger when average direction was towards the right (10°) and the following finger when average direction was towards the left (−10°) in all cases. Grey small points represent mean weights calculated for each participant, whereas big points represent the group-level means. Error bars represent standard error across participants. *represent p<.05/3 between average direction of −10 and 10 degrees. For clarity, weights have been pooled across discrepancy levels and showed across average directions.

Whilst the above analysis showed that finger priority was linked to the average motion direction, we did not assume that the neural process, which determined the weights, involved the estimation of the average direction. It is also unlikely that the weight assignment depended on the process of specifying the VLF because this essentially requires estimating the average angle. Furthermore, the above analysis and results of single-finger stimulations suggested that estimates of component directions could be biased. We, therefore, designed mathematical models that estimate the average direction by integrating distorted estimates of the component directions without explicitly specifying the VLF. One such model had a sensory weighting that depended on the component directions (*biased integration model*) and was capable of prioritising the VLF. This model was compared with an alternative model that assumed equal weights among fingers (*unbiased integration model*). Details of both models can be found in Methods. Briefly, in both models, estimates for each component direction were calculated considering the sensitivity and bias linked to the individual fingers (equation 6). Then, an estimate of the average angle was given as a weighted average of the individual estimates (equations 7 and 8). The biased model included a *gain factor* for each finger and the condition that determined the *strength* to which the finger attracted the weight (equation 10) depending on the direction of stimulus delivered to that finger. Fitting the model and analysing its gain factors or weights allowed us to characterise the direction-dependencies of the sensory weighting that differed across contexts (unimanual vs. bimanual). Fig. 9A shows an example of sensory weighting of the biased model. Fig. 9B shows a possible neural implementation of the biased model based on the normalization model of attention (Reynolds and Heeger 2009).

**Figure 9.**
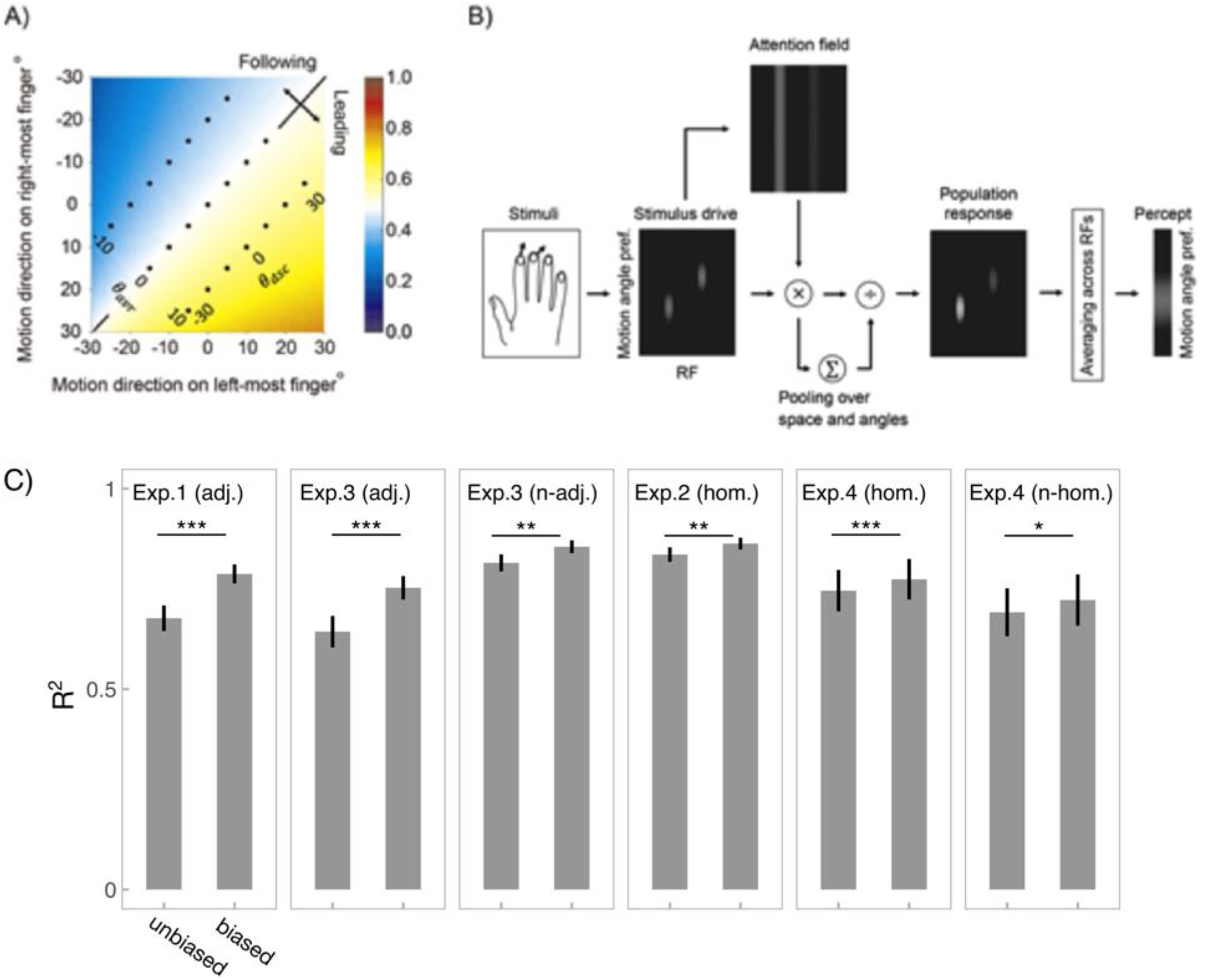
Biased integration model. **A)** Weight assignment based on the biased integration model with gain factors that suffice the virtual leading-finger priority conditions (c1 = 0.01, c2 = −0.01; conditions shown in S5). The colors in the figure show the weight assigned to the left-most finger. Note that the weight is higher in the bottom-right half where the left-most finger would be the virtual leading finger. Black dots show the stimulus pairs used in the experiment. Dots aligned from bottom left to top right have the same average angles, whereas dots aligned from bottom right to top left have the same discrepancies of angles. X and Y axes are aligned to Figure 1C. **B)** An interpretation of the model based on the normalization model of attention (Reynolds and Heeger, 2009). Stimuli on fingers cause initial sensory response (stimulus drive). This modulates the attention gain on each stimulus shown as the attention field. The final neural population response to the stimulus results from divisive normalization of the attention-modulated sensory drive. The percept reflects the biased and normalized population response averaged across the receptive fields. **C)** Coefficient of determination (R^2^) for the two models (biased and unbiased integration model) in the different experiments. Error bars denote standard errors across participants. *, ** and *** represent p<.05, p<.01 and p<.001, respectively.

### Model performance

In the unimanual experiments, the biased integration model provided a better fit for the data (judged angles) compared to the unbiased model (see Table 2). In contrast, there was no such difference in bimanual experiments. Coefficient of determination (R^2^) shown in Fig.9C was significantly higher for the biased integration model in all datasets, but the difference was larger in the unimanual experiments (independent t-tests between Exp1 & Exp2: t_28_ = 2.55, p = .017, d = 0.93; between Exp3(Adj.) & Exp4(Hom.): t_28_ = 3.05, p = .005, d = 1.11; between Exp3(Non-adj.) & Exp4(Non-hom.): t_28_ = 2.51, p = .02, d = 0.92; see S6 for prediction of judgements based on biased model).

**Table 2.**
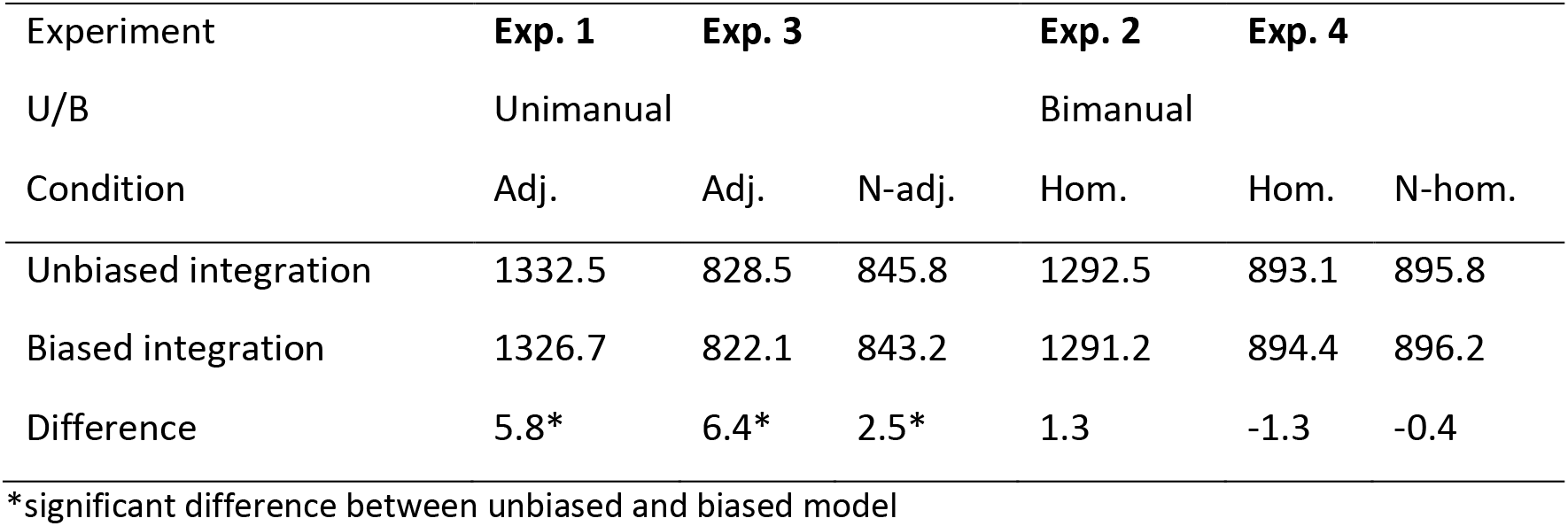
Summary of AIC values (mean value and difference of mean). Significant differences in AIC values are shown in bold letters.

| Experiment | Exp. 1 | Exp. 3 |  | Exp. 2 | Exp. 4 |  |
| --- | --- | --- | --- | --- | --- | --- |
| U/B | Unimanual |  |  | Bimanual |  |  |
| Condition | Adj. | Adj. | N-adj. | Hom. | Hom. | N-hom. |
| Unbiased integration | 1332.5 | 828.5 | 845.8 | 1292.5 | 893.1 | 895.8 |
| Biased integration | 1326.7 | 822.1 | 843.2 | 1291.2 | 894.4 | 896.2 |
| Difference | 5.8* | 6.4* | 2.5* | 1.3 | -1.3 | -0.4 |

\*significant difference between unbiased and biased model

To further confirm that the biased integration model captures participants’ performance, we examined whether it explained the discrepancy-dependent changes in sensitivity to the average direction (see behavioral results). Specifically, if participants averaged component directions with equal weights, the slope would always be unity, independent of the discrepancy. Conversely, the VLF-priority hypothesis predicts a systematic decrease of slope as discrepancies become more positive (i.e., component directions start to *diverge* to the outer edges of the fingerpads). This is because, when component directions *converge* (discrepancy is negative), the direction on the VLF has an equal or larger tilt compared to that on the other finger (see Fig. 10A). Therefore, assigning more weight to the VLF would result in the overestimation of positive angles and underestimation of negative angles, leading to a slope values that exceed 1. When the component directions *diverge* (discrepancy is positive), the predication is opposite, and the slope values decrease. Indeed, when stimuli were delivered to the fingers of the same hand, biased integration model successfully fit the data and showed the monotonic decrease in sensitivity as the discrepancy became more positive (Figs. 10B and 10C). Interestingly, the model also fit the nonlinear trend when discrepancies were negative (i.e., increase in slope values saturated as the discrepancy between component directions decreased). No such discrepancy-dependent modulations were observed in bimanual experiments (Figs. 10D and 10E).

**Figure 10.**
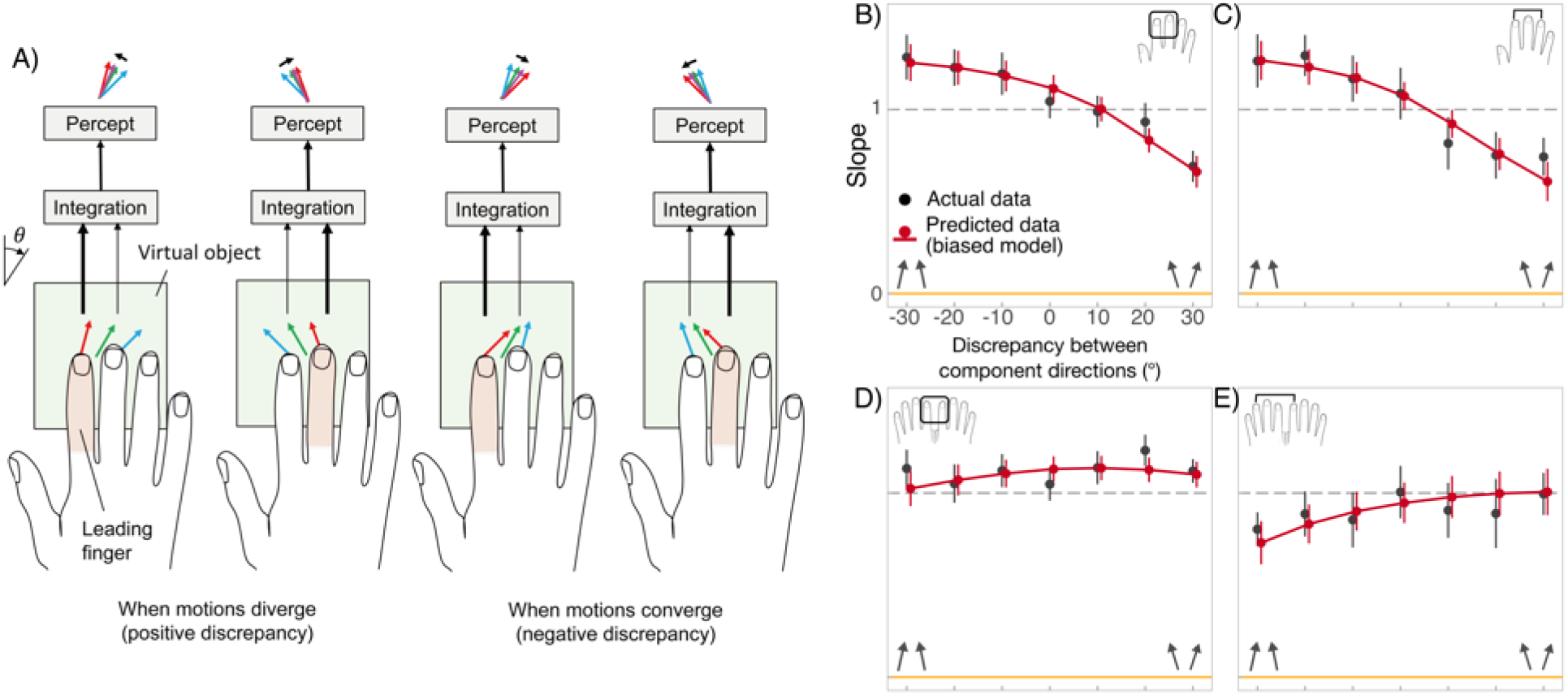
Relation between sensitivity and component angular discrepancy. **A)** How VLF-theory explains the effect of angular discrepancy. Red and blue arrows indicate motions delivered to the *leading* and *following* fingers, respectively. The virtual leading finger is defined by assuming a virtual object (shown as a green box) moving in the direction of the averaged motion direction (green arrow). The theory assumes that the VLF would have a larger influence (indicated as thicker arrows) on perception than the non-leading finger. Importantly, when the discrepancy of the motion direction between the two stimuli is positive (left panel), the motion on the virtual leading finger would have an equal or smaller absolute angle (|θi|) than that of the other finger. Therefore, the theory predicts that the perception (shown as purple arrow on top) would underestimate the size of the average angle (|θavr|). When the discrepancy is negative (right panel), the size would be overestimated. Thus, the VLF-hypothesis predicts that the sensitivity to the average angle would be larger when the discrepancy is negative. Panels **B, C, D,** and **E** show actual data (black dots) and model fit (red dots and overlays) to average angle slope as a function of discrepancy. Dots show group-level slope and error bar stands for standard error across participants. The red overlays show group-level interpolation of biased model predictions. Panel **B** compromises data from unimanual adjacent conditions (N = 30) and panel **C** represents data from non-adjacent condition (N = 15). Panel **D** contains data from bimanual homologous condition (N = 30) and panel **E** represents data from non-homologous condition (N = 15).

### Weight assignment of the biased integration model

After confirming that the biased integration model explained the data, we analysed its weight assignment. Specifically, if VLF is prioritised in the unimanual averaging, gain factors should average to 0 and have a positive difference (see S5 for detail). We confirmed that in all datasets, the average of the gain factors was not significantly different from zero (paired t-tests; adjacent unimanual: t_29_ = .10, p = .92, d = .02; non-adjacent unimanual: t_14_ = .79, p = .45, d = .2; homologous bimanual: t_29_ = 1.53, p = .13, d = .28; non-homologous bimanual: t_14_ = 1.0, p = .33, d = .26). Meanwhile, a significant difference between the gain factors, consistent with the VLF-priority hypothesis, was found both when unimanual finger were adjacent (paired t-test: t_29_ =3.45, p = .002, d = .63) and non-adjacent (t_14_ = 3.96, p = .001, d = 1.0), but not when stimuli were delivered to bimanual fingers, neither when they were homologous (t_29_ = −1.21, p = .24, d = .22) and non-homologous (t_14_ = .84, p = .41, d = .22). Weight assignment calculated from the estimated gain factors for the unimanual conditions are shown in Figure S7. Importantly, direction analysis of the model weights also supported the VLF-hypothesis (see S7 for details).

### Weighting model does not account for average estimate precision

We then examined whether the sensory weighting model could also predict the lower precision when averaging unimanually presented stimuli (see S8 for full details). Since precision of integrated estimate decreases as the weights become biased (Fig. S8.1), the model did predict lower precision in unimanual condition. The question was whether this quantitatively explained the data. We predicted the unbiased SDs in double-finger conditions relative to those in single-finger conditions from the weights of the optimised models. This suggested that the ratio between SDs of unimanual and bimanual averaging would be 1.04 at the largest (when noise in individual estimates is completely independent). The actual ratio (1.09 / 0.91 = 1.2, see Fig. 7) was larger. This suggested that the difference in precision between the unimanual and bimanual experiments cannot be explained solely on the bias of the sensory weighting. Moreover, the degree of sensitivity reduction between converging vs. diverging component directions in unimanual experiments did not correlate with the difference in precision between average vs. component direction estimation (in Exp. 1: *r* = .44, *p* = .10; in Exp. 3 (adjacent): *r* = .002, *p* = 1.0; in Exp. 3 (non-adjacent): *r* = −.15, *p* = .58). Namely, there is no evidence for a shared factor underlying the decrease in sensitivity and in precision under unimanual averaging.

## Discussion

The spatial aspect of touch has remained relatively unexplored – particularly if we set aside the large and productive literature on active touch where motor and proprioceptive signals make a major contribution. Across four experiments, we characterised participants’ ability to aggregate spatiotemporal tactile information across multiple different fingers. Our results show strong integration between multiple tactile inputs, but subject to limitations for inputs delivered within a hand. Our model suggests that tactile inputs are weighted according to an integrative model of hand-object interaction that operates within-hands on purely geometric information to prioritise ‘novel’ information from a ‘virtually leading finger’ (VLF).

### Sensitivity

Several factors may affect sensitivity of tactile direction judgements. For example, perceptual repulsion away from canonical directions has been reported for visual orientation judgements (Jazayeri and Movshon, 2007). A similar effect may occur in touch, potentially explaining the above-unity slopes for judgements of directions on an individual digit. Yet, it may not be an appropriate explanation for judgments of *average* direction between multi-digit stimuli. While overall sensitivity to tactile direction was equal between double- and mean of single-finger conditions, we observed a striking deterioration of sensitivity to *average* direction when component directions diverged from one another. This was found only when those were delivered to the fingers of the same hand (Exp.1 & Exp.3), and not when they were delivered to fingers on different hands (Exp.2 & Exp.4). Our biased integration model successfully predicted the discrepancy-dependent change in sensitivity during unimanual averaging. This is because VLF-priority predicts higher weighting of VLF. Thus, when component directions diverged, the VLF had either equal or smaller absolute angle compared to that on the other finger, resulting in underestimation of the tilt of the average motion.

### Sensory weighting

In support to the VLF-priority hypothesis, we also found that the relative weight assigned to the left-most finger during unimanual averaging shifted according to the average motion angle. No such modulation was found for bimanual averaging. Indeed, while bimanual average direction judgements were equally well predicted by biased and unbiased (assuming equal weights among fingers) integration models, unimanual average judgements were significantly better predicted by the biased integration model. Moreover, the weighting assignment by the model followed the pattern of results predicted by the VLF-priority hypothesis, but only in the unimanual case.

Why does VLF-priority apply to unimanual but not to bimanual averaging? In everyday experience we tend to touch a single object with one hand, but different objects with different hands. Thus, tactile stimuli on different fingers of the same hand are often similar. Predictive coding accounts and other Bayesian approaches show that priors (that reflect the natural statistics of the stimuli) are combined with sensory evidence during the estimation process (Knill and Pouget, 2004; Weiss et al., 2002; Stocker and Simoncelli, 2006). Thus, participants’ behavior in our perceptual averaging task could be explained based on their pre-existing priors (or beliefs) about the statistics of the natural stimuli, particularly for coherent object motion across the multiple fingers of one hand. For such coherent object motions, one finger can be designated as the VLF, and the others as redundant. In our experimental setup, the stimuli were in fact two probes that moved across the static fingertips, in independent directions. The stimulus directions were generally discrepant, and thus incompatible with a coherent object motion. Nevertheless, the distinctive VLF bias associated with coherent object motion was strongly present in the data. This suggests that participants cannot avoid averaging the stimuli in a way that expresses a coherent object motion prior. Many hand-object interactions involve tactile stimulation that is consistent across several digits. However, limited capacity means that not all of this tactile information can be processed. Further, high redundancy means that not all of the tactile information *needs* to be processed. We suggest that an ‘object-motion’ prior offers a useful cognitive heuristic to overcome these limitations, and to optimise tactile integration in typical unimanual touch. Specifically, the VLF provides the least redundant, and therefore most informative tactile signals for guiding the interaction. However, in our task, the multiple tactile signals could have discrepant component directions. Applying an object-motion prior in such cases produced the slightly biased, suboptimal weighting that we observed. In contrast, tactile motions experienced by the two hands are often independent, even during coordinated bimanual actions. This may explain the lack of prioritisation in the bimanual case.

A criticism of our weight analysis could be that when individual estimates are biased, weights can appear biased, when they are, in fact, equal (see S4 for detail). Indeed, discrepancy did affect weighting in the unimanual tasks and in some conditions, we observed significant biases in direction estimation. However, even when each finger has different sensitivities and biases, they cannot explain the difference in the weighting shown in Fig. 8. This is because the effects of the biases are cancelled out when the apparent weight is averaged across the conditions and the difference in the sensitivities only produces fixed shifts in the weights (see S9 for detail). Another criticism could be that there might be other forms of biased sensory weighting. We addressed this concern by comparing our model with numerous alternative models, including models that omitted components of the biased integration model and models that assumed lateral interaction among fingers (see S10). Importantly, we found that the models with lateral-interaction showed worse fit to the judgements compared to the biased-integration model. This is consistent with the finding that the VLF-priority hypothesis extended to a case with non-adjacent fingers, and in general, somatotopic distance between stimuli did not appear to modulate averaging (Exp.3).

How does weighting of tactile input occur in real time? Because our stimuli lasted approximately 1 s, the weighting had to be applied within that short time window. Although, stimulus-driven shifts in tactile attention occur rapidly (in less than 200 ms; Spence and McGlone, 2001), the computational processes involved in VLF-priority may seem more demanding than a simple exogenous shift of attention. Intuitively, the brain must first estimate the motion direction of an implicit virtual object moving across the skin. Next, the brain must identify the VLF given the implied object motion. Finally, the increased weighting to prioritise input from the VLF must be implemented, perhaps by a shift of attention. By contrast, we show that a simple network can automatically prioritise the VLF without estimating the object motion and without explicitly identifying the VLF. In the model, each stimulus attracts attentional resources with a strength that depends on its direction and the fingers stimulated. The process is optimised so that stimulus that appears to be on the VLF is more salient. This may explain why the computation can be performed in real-time, thus contributing to perception.

Our model extends on existing models of tactile perception. For example, Pei et al. (2011) showed that perception of tactile plaid motions depends on a saliency-dependent weighting of each motion component. Although our model was implemented as an angle summation model, it is approximately equal to a vector summation model (see S11). While saliency-based integration theories (Cataldo et al., 2019; Walsh et al., 2016) and other reliability-based integration frameworks (Ernst and Banks, 2002) explain general biases in integration, they fail to explain the discrepancy effect, which results in biases even when salience and reliability is held constant. We show that discrepancy effect can be explained by assuming the VLF-priority. Thus, our finding suggests a novel type of sensory weighting that depends on the expected novelty of each stimulus.

### Precision

While the model successfully explained the averaging performance in terms of sensitivity, another major finding was that averaging directions bimanually led to increased precision as compared to estimating each direction in isolation. This precision benefit did not occur for unimanual stimuli. By applying our biased weighting model to unbiased direction estimate SDs, we found that additional factors may be required to explain the precision difference.

Results of Experiments 3 and 4 lead us to reject that the effect is due to somatotopically organised inter-digit interactions. Although caution is required in drawing conclusions from null results, our findings are unlikely to simply reflect lack of power as Bayesian analysis tended to support the null effects. In addition, Walsh et al. (2016) also failed to observe somatotopic distance effects during unimanual aggregation of intensity-based tactile signals. This is also consistent with our previous findings (Arslanova et al., 2021) showing that suppressive interactions between digits may be reduced when participants try to estimate average directions, as opposed to direction discrepancies. Thus, somatotopic interactions might have been attenuated in our averaging conditions. The absence of somatotopic effects could be related to the level of abstraction that is involved in perceiving an ensemble property across spatially distinct inputs (for ensemble perception in vision see Alvarez, 2011; Whitney and Yamanashi Leib, 2018). Thus, perceiving an average direction between component trajectories may no longer depend on somatotopically organised circuitry. The ability to abstract sensory information away from the specific receptive fields stimulated is an important precursor for object and event perception (Marr, 2010).

An alternative explanation is based on constrained attentional resources that may have prevented the parallel processing of concurrently delivered trajectories to a single hand, resulting in suboptimal perception of average direction. Our weighting results showed biased weighting during unimanual averaging, but they did not find total selectivity, in which signals from one finger would be entirely ignored. This suggests that information from both fingers was processed. However, the reduction in precision during unimanual averaging compared to estimating each component in isolation suggests that the two signals were compromised when presented simultaneously. The improved relative precision in bimanual task implies improved ability to process the two signals in parallel. This could be due to the engagement of two separate pools of processing resources corresponding to each hand. Although some authors find performance costs for bimanual tasks (Nguyen et al., 2020), the idea that perception can benefit from distributing resources across hemispheres is well documented (Friedman and Polsen, 1981; Schweinberger et al., 1994; Bradshaw et al., 1998). For example, Craig (1985) and Overvliet et al., (2007) both showed that participants were better at identifying tactile patterns when the pattern components were presented bimanually rather than unimanually.

Yet another explanation relies on theories of noise summation, whereby when multiple signals are each affected by an independent source of noise, averaging those signals leads to increased precision (Zohary et al., 1994; Averbeck et al., 2006; Alvarez, 2011). Accordingly, precision differences in our tasks may have arisen because multiple stimuli encoded within the same hand were affected by shared sensory noise, whereas same stimuli encoded on different hands were affected by relatively independent noise. Consistently, Cohen and Maunsell (2009) found that variability in individual neurons’ firing rates correlated within a hemisphere, indicating a shared variability throughout the neural population, whereas noise correlations in different hemispheres were close to zero, suggesting that sensory noise between the two hemispheres remains independent. This is consistent with our finding that precision was not modulated by somatotopic distance between component directions. A potential source of such noise interdependence for stimuli encoded within the same hand could be fluctuations in the mono-hemispheric somatosensory rhythms (Andrew and Pfurtscheller, 1999; Kilner et al., 2003).

Since our study was behavioural, we cannot draw strong conclusions regarding the neural basis of our participants’ percepts. Specifically, we cannot measure whether our bimanual averaging task engaged stronger inter-hemispheric processing, given that unimanual stimuli are also known to engage both hemispheres (Tamè et al., 2016; Iwamura et al. 2002). Yet, the ipsilateral activation following unimanual stimulations is lower than its contralateral counterpart, and presumably arises via transcallosal input from the contralateral SI (Sutherland and Tang, 2006). Further, bilateral activation would always be stronger, and more symmetric, for bimanual than for unimanual stimuli. Together with the finding that bimanual averaging was not affected in a digit-specific manner (Exp.4), we believe that differences between averaging unimanual and bimanual stimuli correspond to largely hand- or hemisphere-specific factor. Additional neuroimaging or EEG methods could confirm the hemispheric lateralisation of our stimuli, and to provide direct support for this conclusion.

Taken together, our work reveals two distinctive features of within-hand and between-hand tactile motion integration. First, we identified a process of sensory weighting that involves a prior belief based on natural tactile interactions, and that operates for the digits within one hand. Specifically, tactile processing incorporates an *implicit* prior consistent with a single object moving across the skin. As a result, during unimanual averaging, the VLF is expected to receive novel information, thus biasing perception. We showed that this biasing occurs, even in tasks like ours, where no novelty actually exists, in the strict temporal sense of the term, and where implicit object motion is known only geometrically. We showed that this biasing effect of ‘natural’ priors provides an accurate prediction of our perceptual averaging data, notably the relation between sensitivity and discrepancy. Future studies might design experiments specifically targeting the theoretical constructs of the VLF-priority theory. For instance, such experiments might specifically manipulate the ‘single object’ prior (e.g., by visually showing either one or two moving objects during tactile stimulation). Second, analysis of precision showed that the capacity to aggregate stimuli within one hand may be limited. This limitation could be due to restricted processing resources or shared noise between fingers of the same hand. As a result, bimanual touch has a perceptual advantage in terms of precision. Based on these results, we argue that the aforementioned ‘object-motion’ prior may serve as a cognitive heuristic to overcome the limited physiological resources for optimal integration in unimanual touch. Ultimately, our findings reveal the power of object-level perception in the context of an afferent system whose basic sensory capacities are relatively limited.

## Methods

### Participants

In total, 60 participants took part in the study, 15 in each of 4 experiments. All participants gave informed consent prior to experiments, in accordance to the declaration of Helsinki. The study was approved by the University College London Research Ethics Committee. Seven participants were excluded from analysis (four in Exp 1; one in Exp 2; one in Exp 3; one in Exp 4), because they had estimation errors exceeding 20 degrees in more than 50% of trials in at least one single-finger condition, making their data unreliable. Excluded participants were replaced with others. The demographics of the final sample were as follows: Experiment 1 (age range: 21 – 39; mean: 27.07; 10 women, 5 men; all but one reportedly right-handed); Experiment 2 (age range: 22 – 40; mean: 25.87; 9 women, 6 men; all reportedly right-handed); Experiment 3 (age range: 18 – 30; mean: 23.47; 11 women, 4 men; all but one reportedly right-handed); Experiment 4 (age range: 18 – 34; mean: 22.20; 10 women, 5 men; all reportedly right-handed). A sample size of 15 was estimated using G*Power 3.1 (Faul et al., 2009), based on desired power of 0.80 and an effect size of 0.68 for aggregated versus single-digit motion perception precision (unpublished pilot study).

### Tactile apparatus

It consisted of two spherical probes (4 mm diameter) attached to two stepper linear actuators (Haydon Kerk Motion Solutions 15000 series, model LC1574W-04) that were fixed to two motorized linear stages (Zaber X-LSM100B, Zaber Technologies Inc., Canada) mounted in an XY configuration (Fig. 1D). The actuators were controlled by a microcontroller (Arduino) and were moved vertically so that a plastic hemispheric probe (diameter 4 mm) could make/break static contact with the fingerpad skin at the start of tactile stimulation and retract after the end of stimulation. The probe could be moved horizontally in predefined trajectories (see *Tactile stimuli* below). The apparatus was covered by a box with a small aperture. To-be-stimulated fingertips were positioned in a palm-down position over the aperture and secured with foam padding. During the stimulation the probes lightly touched the fingerpads. A webcam was placed under the apparatus to monitor the finger placement and contact with the probe. The aperture was then covered with a screen, so that participants could not see the stimuli. The same apparatus was used for all experiments.

### Tactile stimuli

Continuous motion along the fingertips was created by moving the probes at preselected angles ranging from −25 to 25 degrees to the distal-proximal finger axis in 5° steps, at a constant speed of 10 mm/s. The movement of each probe was controlled individually allowing for delivery of trajectories with varying discrepancy simultaneously along both fingertips. Figure 1C shows 21 possible combinations compromised of 11 individual directions delivered simultaneously to two fingers. The combinations produced three different average motion patterns, with varying discrepancy between the two stimuli. The sign of discrepancy reflects whether the component directions tended to converge towards the inner edges of fingertips (negative discrepancy) or diverge towards the outer edges of fingertips (positive discrepancy). At the beginning of each trial, the probe was advanced to make a static contact with the fingertip. The initial position of the probe was jittered across trials (−2.0, 0.0, or 2.0 mm from the center of the fingertip) to discourage using memory for locations as a proxy for direction. After each trajectory, the probe was immediately retracted and returned to its starting position. The duration of each trajectory was approximately 1 s and the distance travelled was 10 mm.

### Experimental design and procedure

In all conditions, participants rested their hand(s) in a fixed palm-down position. In Experiment 1, the probes, through the aperture, lightly touched the centre of their right index and middle fingertips. In Experiment 2, the probes touched the right index and left index fingertips. In Experiment 3, the set-up was identical to Experiment 1, and additionally included a non-adjacent condition, in which the probes touched the right index and right ring fingers. In Experiment 4, the set-up was identical to Experiment 2, and additionally included a non-homologous condition, in which the probes touched the left index and right ring fingers. The distance between fingers and corresponding width between probes was held fixed across conditions whenever possible in order to minimise the effects of spatial distance between fingers. Thus, in Experiment 1 and 2, the distance between fingers was fixed to approximately 25 mm. In Experiment 3, fingers in the adjacent condition remained at ~25 mm; in the non-adjacent condition, the distance between index and ring finger was ~45 mm. In Experiment 4, the distance between bimanual fingers was fixed to ~65 mm in order to make homologous and non-homologous conditions comparable without the confounding effects of spatial separation.

To investigate perception of the average motion pattern from two separate trajectories, we compared perception of the average direction to perception of the two individual component stimuli presented alone. Accordingly, all experiments contrasted double-finger stimulations with the single-finger stimulations of which they were composed. Single-finger conditions were repeated for each finger (e.g., in Experiment 1 participants performed estimation on index finger and separately on middle finger). The single-finger conditions for each finger were then averaged to obtain one measure characterising single-finger perception. In all conditions, participants gave their response after the motion stimuli ended, by adjusting the orientation of a visual arrow that appeared on the computer screen placed immediately above their fingertips. They had to adjust the arrow’s orientation to the perceived single-finger direction (single-finger condition) or average direction (double-finger condition). The adjustment was made either by pressing left and right arrow keys (and enter key to record their final response) with their (unstimulated) left hand in Experiments 1 and 3, or by pressing anticlockwise and clockwise rotator pedals with the left foot (and a third pedal with the right foot to record their response) in Experiments 2 and 4. Responses were unspeeded, and no feedback was given. After the response, the arrow disappeared, and the probes moved to their starting positions.

In Experiments 3 and 4, we further examined how averaging ability changes as a function of somatotopic distance between fingers. We compared the contrast between estimating individual directions (single-finger condition) and estimating their average (double-finger condition) across two different somatotopic distances. In Experiment 3 the fingers were either adjacent (right index and right middle) or non-adjacent (right index and right ring). In Experiment 4 the fingers stimulated were either bimanually homologous (right index and left index) or non-homologous (right ring, left index).

All conditions were blocked. In Experiments 1 and 2, the order of finger conditions was counterbalanced across participants. In Experiments 3 and 4, the order of somatotopy conditions (adjacency or homology) was counterbalanced across participants, and finger condition was counterbalanced within each somatotopy condition. In double-finger conditions, the 21 stimulus combinations were repeated 7 times in Experiment 1 and Experiment 2, 5 times in Experiment 3 and 6 times in Experiment 4. In single-finger conditions, the number of repetitions of each direction was matched to that in the double-finger condition. The total number of trials was 168 per condition in Experiment 1 and Experiment 2, 105 per condition in Experiment 3 and 120 per condition in Experiment 4. Experiments 1 and 2 lasted approximately 1.5 hours, while Experiments 3 and 4 lasted approximately 2 hours and were performed across two 1-hour sessions on separate days.

### Analysis of main measures: sensitivity, bias, and precision

Slope and intercept values were estimated by fitting linear regressions to each participant’s data separately in each condition. Group-level adjusted r^2^ for linear fits for single-finger conditions were as follows: .61 ± .07 in Exp1, .57 ± .11 in Exp2, .65 ± .09 in Exp3 (adjacent), 62 ± .09 in Exp3 (non-adjacent), .57 ± .15 in Exp4 (homologous), and .55 ± .17 in Exp4 (non-homologous). For double-finger conditions: .28 ± .12 in Exp1, .42 ± .11 in Exp2, .34 ± .11 in Exp3 (adj), .28 ± .13 in Exp3 (non-adj), .42 ± .17 in Exp4 (hom), and .37 ± .18 in Exp4 (non-hom).

Unbiased SDs were measured over repeated *different* tactile motion stimuli. In other words, in single-finger conditions, SDs were measured for each angle. Some angle-specific effects were present in single-finger conditions (see S1), but these were minimal. We took the mean of angle SDs to reflect the inverse precision for single-finger condition. In double-finger condition, SDs were measured for each combination. There were no combination-specific effects (see Table S1). The mean of this value reflects inverse precision for double-finger condition. Because we wanted to keep the stimuli in single-finger conditions identical to the ones in double-finger conditions, the number of trials for each single-finger angle varied (i.e., 0° was used 28 times, whereas 25° only 7 times) to match the number of times each was used in double-finger combinations. However, normal sample SD is a biased estimate of the population SD – the smaller the sample size the more likely it will underestimate the population SD (Montgomery and Runger, 2010, section 7.3). Therefore, to account for different number of trials in each angle, we used an unbiased SD to calculate precision over angles and combinations with the following equation:

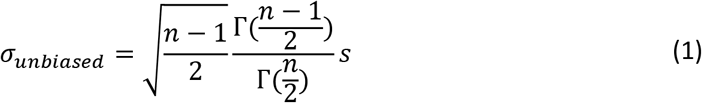

where Γ(·) is the Gamma function and s represents the usual standard deviation.

In all experiments, we first used one-sample t-tests on slope and intercept values to see whether participants could perceive probes’ directions and whether their perception was unbiased. When normality was violated, we used the sign test instead. In Experiments 1 and 2, we were interested whether the aggregation process differs according to whether component directions are combined within- or between-hands. We carried out a mixed ANOVA with within-subjects factor number of fingers (single-finger vs. double-finger condition) and between-subject factor experiment (Experiment 1: unimanual vs. Experiment 2: bimanual) on sensitivity, bias and precision. In Experiment 3, the main interest was whether within-hand averaging ability depended on the somatotopic distance between fingers. A repeated-measures ANOVA was employed with factors number of fingers (single-finger vs. double-finger condition) and adjacency (adjacent vs. non-adjacent fingers) on the three measures. In Experiment 4, the main interest was whether the putative somatotopic mechanism extends to between-hands averaging. Here, similarly to Experiment 3, repeated-measures ANOVA were employed with factors number of fingers (single-finger vs. double-finger condition) and homology (homologous vs. non-homologous fingers).

For linear models such as ANOVA, normality assumption should be checked not against raw dependent variable, but on the residuals (or errors) from the fitted model (Kozak and Piepho, 2017). However, ANOVA with a balanced design is considered robust even with normality violations, thus, violations of parametric assumptions were unlikely to majorly influence Type I and Type II errors (Boneau, 1960; Glass et al., 1972). In addition, alternative non-parametric Friedman test cannot be done for factorial designs. For these reasons, parametric ANOVA was used throughout.

In addition to the main analysis, we explored whether averaging ability varied as a function of discrepancy between the component directions. Therefore, we fit separate linear regressions to the double-finger data for each discrepancy level and extracted the slope and intercept values for each discrepancy. Discrepancy of 0, when component directions were identical, was excluded from the analyses, as those stimuli did not have two signs of direction. For precision, unbiased SDs were calculated to each pair of stimuli (3 average directions by 6 levels of discrepancy). For Experiments 1 and 2, mixed ANOVA with within-subjects factors sign of discrepancy (diverging vs. converging) and level of discrepancy (30° vs. 20° vs. 10°) and between-subjects factor experiment (unimanual vs. bimanual) was carried out. For Experiment 3, repeated-measures ANOVA was employed with factors sign of discrepancy (diverging vs. converging), level of discrepancy (30° vs. 20° vs. 10°), and adjacency (adjacent vs. non-adjacent fingers). For Experiment 4, the same analysis was carried out, except adjacency factor was replaced with homology (homologous vs. non-homologous fingers). When modeling precision, average direction was added as an additional within-subject factor to reveal any direction-specific differences.

Lastly, support for the null hypothesis could be scientifically informative, as well as support for the experimental hypothesis. Therefore, we used Bayes Factors (Jeffreys, 1961; Rouder et al., 2012; Wagenmaker et al., 2018a, 2018b) to assess support for the null hypotheses, in appropriate cases. BF^01^ was used to indicate the strength of evidence for the absence of an effect/interaction over a model that did not contain that specific effect/interaction, while BF_inclusion_ was used to reflect the strength of evidence for including a particular effect/interaction against all other models. Following the guidelines by Kass and Raftery (1995) we considered BF > 3 and BF < 0.33 showing sufficient evidence, while 0.33 < BF < 3 showing inconclusive evidence.

The data used for all experiments, the main estimated measures, and analysis scripts are openly accessible on Open Science Framework (https://osf.io/26ng3/?view_only=025924e13fbb4cf8b75da20d1b9709ae).

### Weight estimation from the direction judgments

In order to infer the relative contribution of each finger in making the direction judgments, we estimated the weight given to the left-most finger (right index finger in unimanual conditions and left index finger in bimanual conditions) using the following equation.

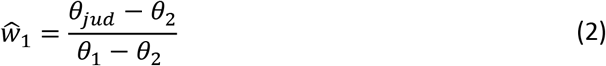

*θ*_1_ and *θ*_2_ represent the angle of motion given to the left- and right-most finger, respectively. *θ*_*jud*_ represents the judged angle. Angle was positive when the tactile motion headed towards the right from the participant’s point of view (i.e., towards the right-hand side; see Figure 1). We only calculated the weight for the left-most finger since the weight for the other finger would simply be 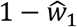. The equation assumed that the judgements were based on weighted averaging of the individual angles as follows.

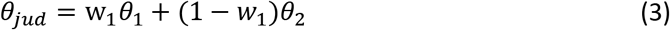

We estimated the weight for all the tested stimuli except for those in which the discrepancy of the angles

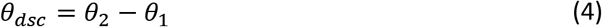

was 0 (when the angles were identical between the two fingers; note that weight calculation is impossible in this case). Since the functionally leading finger should switch between conditions in which the average angle

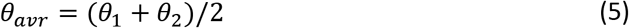

Since the VLF-priority hypothesis predicted that the weight would depend on the average angle (main effect) and would be higher when the average angle is 10° (when the finger leads) compared to when it is −10° (when the finger follows). For that, we applied four separate ANOVAs to the estimated weights in the four conditions: adjacent unimanual, non-adjacent unimanual, homologous bimanual, and non-homologous bimanual. In each case, the within-subject factors were average direction (−10°, 0°, 10°), direction of discrepancy (diverging, converging) and level of discrepancy (30°, 20°, 10°). In the adjacent and homologous conditions, data was merged across two experiments. Therefore, experiment was added as a between-subject factor in these conditions.

### Sensory weighting model

We designed a mathematical model of weight assignment that can prioritize the leading finger without estimating the average angle or the leading finger (*biased integration model*, described below). The model was compared with an alternative model that assumes equal weights among fingers (*unbiased integration model*). We used iterative least squares estimation to fit both models to the judgements. A single set of parameters was used to predict all the judgements made by the same participant within the same somatotopic condition. The models were compared based on the Akaike information criteria (AIC). As is conventional, difference in mean AIC was considered meaningful when it exceeded 2.0 (Burnham and Anderson 2004). The design of the biased integration model was chosen by comparing various alternative models using the same criterion (see S10 for detail).

Both biased and unbiased models assume that each finger provides an estimate of the motion angle in the following form

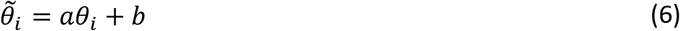

and that the estimate of the average angle 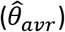 is given as a weighted average of the individual estimates 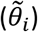 as shown below.

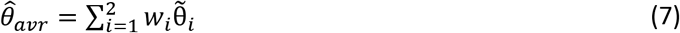

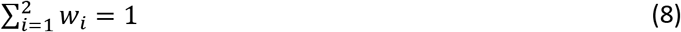

The index i=1,2 corresponds to the left-most and left-hand-side fingers, respectively. Equation (6) was based on the result of single finger stimulation which suggested that the judgement can be fitted by such a linear model. The formalization considers the possibility that the individual estimates are biased. While it was possible that the two fingers have different sensitivities (a) and biases (b), we assumed that the two values were shared among the two fingers based on three pieces of evidence. First, assuming different values for each finger made the optimization process unstable and did not allow a reliable estimation of the model parameters. Secondly, using the sensitivities and biases identified for each finger by fitting the judgments made in single finger experiments did not improve the fitting performance (see S10 for detail). Finally, our model, which assumed shared sensitivities and biases, accurately predicted the judgements.

Equations (7) and (8) are equivalent to equation (3) except for the fact that the integrated information is now based on biased estimates of the individual fingers. Details on each model of weight allocation are explained below.

### Unbiased-integration model

This model assumes that weights are allocated equally to the two fingers (i.e.,*w*_1_ = *w*_2_ = 1/2). The final estimate would thus be as follows

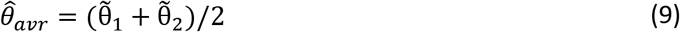

The model had two parameters (*a*, *b* in equation 5) and the parameters were determined by fitting the model to the judgements.

### Biased-integration model

This model assumes that the strength (*g_i_*) in which each finger attracts the weight depends on the angle of tactile motion delivered to that finger as follows.

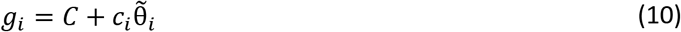

*c_i_* (i=1,2) are finger- and context-specific gain factors which allow the different fingers to have different direction-dependencies in different contexts (unimanual/bimanual). *C* is a constant which can be any positive real number (in our analysis, it was set to 1/2 to ease interpretation). The weights allocated to the fingers are calculated by normalizing the strengths as follows.

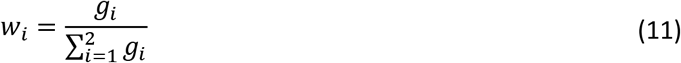

The weights were constrained to have a value between 0 and 1. The model had four parameters in total (*a*, *b*, *c*_1_, *c*_2_) and the parameters were determined by fitting the model to the judgements.

The model can assign the weights to the fingers in various ways depending on the gain factors. When there are two fingers, as was the case in our study, the condition for the biased integration model to always assign more weight to the leading finger (defined by average angle) is given as follows (see S5 for details).

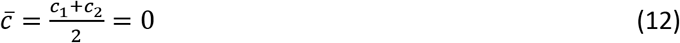

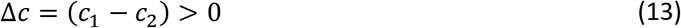

Our model only specifies a mathematical relation between the motions and the judgements. However, a neural implementation of the model could be considered based on the normalization model of attention (Reynolds and Heeger, 2009) as follows (see also Fig. 9B). The stimuli delivered to the two fingers cause the initial sensory responses (stimulus drive) that correspond to 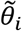 in eq. 6. The saliencies of each stimulus depend both on the motion directions and the context. This results in different levels of attentional gain on each stimulus, the gain represented as *g_i_* in eq. 10 and shown as the attention field in the figure. The final population response to the stimulus results from divisive normalization (eq. 11) of the attention-modulated sensory drive. The percept reflects the biased and normalized population response averaged across the receptive fields.

## Acknowledgments

This study was supported by a research contract between NTT and UCL, and an MRC-CASE studentship MR/P015778/1. We thank Gertheka Sritharan for helping collect the data for Experiment 4.

