## Supplemental information for "Sensory and cognitive factors affecting multi-digit touch: a perceptual and modeling study"

**Contents**

S1. Full statistical analysis of aggregation performance

S2. Single-finger conditions

S3. Full statistical analysis of weighting data derived from data and model

S4. Effect of judgement biases on the estimation of weights

S5. Condition for prioritising the leading finger in the biased integration model

S6. Judgement prediction of the biased-integration model

S7. Weight assignment of the model

S8. Relation between biased sensory weighting and decrease in precision

S9. Difficulty to explain the effect of average angle on weights based on sensitivities and biases

S10. Alternative models for the virtual-leading-finger priority (VLF-priority)

S11. Weighted averaging of vectors and weighted averaging of angles

**S1: Full statistical analysis of aggregation performance**

**Table S1**

Full statistical analysis of aggregation performance

| | | | |
|---|---|---|---|
| SENSITIVITY | | | |
| **Experiments 1 and 2** single- vs. double- condition: | | | |
| Experiment (Exp.1 vs. Exp.2; between) | $F_{1, 28}$ = .90, $p$ = .35, $\eta p^2$ = .03 | $BF^{01}$ = 2.0 ± 1.7% $BF_{inclusion}$ = .5 | |
| Condition (single vs. double; within) | $F_{1, 28}$ = 3.5, $p$ = .08, $\eta p^2$ = .11 | $BF^{01}$ = .5 ± 1.4% $BF_{inclusion}$ = .9 | |
| Experiment x Condition | $F_{1, 28}$ = 2.24, $p$ = .15, $\eta p^2$ = .07 | $BF^{01}$ = 1.2 ± 2.3% $BF_{inclusion}$ = .6 | |
| **Experiment 3** single- vs. double-condition: | | | |
| Adjacency (within) | $F_{1, 14}$ = .45, $p$ = .51, $\eta p^2$ = .03 | $BF^{01}$ = 3.5 ± 1.0% $BF_{inclusion}$ = .3 | |
| Condition (single vs. double; within) | $F_{1, 14}$ = 16.5, $p$ = .001, $\eta p^2$ = .54 | $BF^{01}$ < .001 ± 2.0% $BF_{inclusion}$ = 984.5 | slope larger in single than double |
| Adjacency x Condition | $F_{1, 14}$ = 1.5, $p$ = .24, $\eta p^2$ = .10 | $BF^{01}$ = 1.9 ± 3.3% $BF_{inclusion}$ = .5 | |
| **Experiment 4** single- vs. double-condition: | | | |
| Homology (within) | $F_{1, 14}$ = 4.3, $p$ = .06, $\eta p^2$ = .23 | $BF^{01}$ = 1.7 ± 1.7% $BF_{inclusion}$ = .5 | |
| Condition (single vs. double; within) | $F_{1, 14}$ = 2.0, $p$ = .18, $\eta p^2$ = .13 | $BF^{01}$ = .9 ± .9% $BF_{inclusion}$ = .9 | |
| Homology x Condition | $F_{1, 14}$ = .70, $p$ = .42, $\eta p^2$ = .002 | $BF^{01}$ = 2.1 ± 9.2% $BF_{inclusion}$ = .4 | |
| **Exp. 1 and 2** averaging as a function of discrepancy: | | | |
| Discrepancy level (within) | $F_{2, 56}$ = .48, $p$ = .62, $\eta p^2$ = .02 | $BF^{01}$ = 13.8 ± 1.7% $BF_{inclusion}$ = .03 | |
| Discrepancy sign (within) | $F_{1, 28}$ = 6.1, $p$ = .02, $\eta p^2$ = .18 | $BF^{01}$ = .42 ± 1.0% $BF_{inclusion}$ = 64.5 | slope larger for converging directions |
| Experiment x Disc. level | $F_{2, 56}$ = .21, $p$ = .81, $\eta p^2$ = .007 | $BF^{01}$ = 9.4 ± 3.7% $BF_{inclusion}$ = .02 | |
| Experiment x Disc. sign | $F_{1, 28}$ = 11.5, $p$ = .002, $\eta p^2$ = .29 | $BF^{01}$ = .01 ± 5.6% $BF_{inclusion}$ = 92.2 | |
| Follow-up (bonferroni paired tests) | effect of sign in Exp.1: $p$ < .001 effect of sign in Exp.2: $p$ = .45 | | larger slope for converging directions only present in unimanual experiment |
| Experiment x Disc. level x Disc. sign | $F_{2, 56}$ = .68, $p$ = .51, $\eta p^2$ = .02 | $BF^{01}$ = 3.5 ± 16.9% $BF_{inclusion}$ = .006 | |
| **Exp. 3** averaging as a function of discrepancy: | | | |
| Discrepancy level (within) | $F_{2, 28}$ = .45, $p$ = .64, $\eta p^2$ = .03 | $BF^{01}$ = 114.8 ± .9% $BF_{inclusion}$ = .04 | |
| Discrepancy sign | $F_{1, 14}$ = 12.7, $p$ = .004, $\eta p^2$ = .48 | $BF^{01}$ < .001 ± 1.2% | slope larger for |

| | | | |
|---|---|---|---|
| (within) | | $BF_{inclusion} > 20000$ | converging directions |
| Adjacency x Disc. level | $F_{2, 28} = .40$, $p = .62$, $\eta p^2 = .03$ | $BF^{01} = 7.2 \pm 6.6\%$ $BF_{inclusion} = .01$ | |
| Adjacency x Disc. sign | $F_{1, 14} = 1.4$, $p = .25$, $\eta p^2 = .09$ | $BF^{01} = 3.1 \pm 4.1\%$ $BF_{inclusion} = .22$ | |
| Adj. x Disc. level x Disc. sign | $F_{2, 28} = .10$, $p = .34$, $\eta p^2 = .06$ | $BF^{01} = 3.0 \pm 4.6\%$ $BF_{inclusion} < .001$ | |
| **Exp. 4** averaging as a function of discrepancy: | | | |
| Discrepancy level (within) | $F_{2, 28} = .08$, $p = .92$, $\eta p^2 = .006$ | $BF^{01} = 16.8 \pm 1.0\%$ $BF_{inclusion} = .02$ | |
| Discrepancy sign (within) | $F_{1, 14} = .81$, $p = .38$, $\eta p^2 = .05$ | $BF^{01} = 3.2 \pm 2.9\%$ $BF_{inclusion} = .13$ | |
| Homology x Disc. level | $F_{2, 28} = .04$, $p = .96$, $\eta p^2 = .002$ | $BF^{01} = 7.8 \pm 15.6\%$ $BF_{inclusion} = .01$ | |
| Homology x Disc. sign | $F_{1, 14} = .06$, $p = .81$, $\eta p^2 = .004$ | $BF^{01} = 4.0 \pm 11.7\%$ $BF_{inclusion} = .08$ | |
| Hom. x Disc. level x Disc. sign | $F_{2, 28} = .80$, $p = .46$, $\eta p^2 = .05$ | $BF^{01} = 3.0 \pm 6.9\%$ $BF_{inclusion} < .001$ | |
| BIAS | | | |
| **Experiments 1 and 2** single- vs. double- condition: | | | |
| Experiment (Exp.1 vs. Exp.2; between) | $F_{1, 28} = 12.1$, $p = .002$, $\eta p^2 = .30$ | $BF^{01} = .02 \pm 1.9\%$ $BF_{inclusion} = 13.7$ | bias in unimanual |
| Condition (single vs. double; within) | $F_{1, 28} = .96$, $p = .34$, $\eta p^2 = .03$ | $BF^{01} = 44.7 \pm 1.8\%$ $BF_{inclusion} = .4$ | |
| Experiment x Condition | $F_{1, 28} = .55$, $p = .46$, $\eta p^2 = .02$ | $BF^{01} = 2.3 \pm 3.7\%$ $BF_{inclusion} = .5$ | |
| **Experiment 3** single- vs. double-condition: | | | |
| Adjacency (within) | $F_{1, 14} = 1.4$, $p = .26$, $\eta p^2 = .08$ | $BF^{01} = 2.0 \pm 1.0\%$ $BF_{inclusion} = .4$ | |
| Condition (single vs. double; within) | $F_{1, 14} = .001$, $p = .97$, $\eta p^2 = .001$ | $BF^{01} = 3.8 \pm 3.0\%$ $BF_{inclusion} = .2$ | |
| Adjacency x Condition | $F_{1, 14} = .16$, $p = .69$, $\eta p^2 = .01$ | $BF^{01} = 2.8 \pm 3.1\%$ $BF_{inclusion} = .1$ | |
| **Experiment 4** single- vs. double-condition: | | | |
| Homology (within) | $F_{1, 14} = 1.6$, $p = .22$, $\eta p^2 = .10$ | $BF^{01} = 1.4 \pm .7\%$ $BF_{inclusion} = .7$ | |
| Condition (single vs. double; within) | $F_{1, 14} = .06$, $p = .80$, $\eta p^2 = .005$ | $BF^{01} = 3.7 \pm 1.4\%$ $BF_{inclusion} = .3$ | |
| Homology x Condition | $F_{1, 14} = 6.45$, $p = .02$, $\eta p^2 = .32$ | $BF^{01} = .5 \pm 2.2\%$ $BF_{inclusion} = .8$ | |
| Follow-up (paired-sample t-tests) | Effect of condition in homologous pair: $p = .01$, $d = .75$; Effect of condition in non-hom. pair: $p = .20$, $d = -.34$ | | Bias in double-finger condition, but only during homologous stimulation |
| **Exp. 1 and 2** averaging as a function of discrepancy: | | | |
| Discrepancy level | $F_{2, 56} = 2.35$, $p = .11$, $\eta p^2 = .08$ | $BF^{01} = 9.4 \pm .9\%$ | |

| | | | |
|---|---|---|---|
| (within) | | $BF_{inclusion} = .06$ | |
| Discrepancy sign (within) | $F_{1,28} = 1.2$, $p = .28$, $\eta p^2 = .07$ | $BF^{01} = 2.8 \pm 1.7\%$<br>$BF_{inclusion} = .92$ | |
| Experiment x Disc. level | $F_{2,56} = 3.1$, $p = .10$, $\eta p^2 = .08$ | $BF^{01} = 4.8 \pm 5.3\%$<br>$BF_{inclusion} = .06$ | |
| Experiment x Disc. sign | $F_{1,28} = 3.9$, $p = .06$, $\eta p^2 = .12$ | $BF^{01} = .21 \pm 4.6\%$<br>$BF_{inclusion} = 3.23$ | |
| Experiment x Disc. level x Disc. sign | $F_{2,56} = 1.3$, $p = .28$, $\eta p^2 = .03$ | $BF^{01} = 4.6 \pm 4.7\%$<br>$BF_{inclusion} = .006$ | |
| **Exp. 3** averaging as a function of discrepancy: | | | |
| Discrepancy level (within) | $F_{2,28} = .40$, $p = .64$, $\eta p^2 = .03$ | $BF^{01} = 13.2 \pm 1.4\%$<br>$BF_{inclusion} = .03$ | |
| Discrepancy sign (within) | $F_{1,14} = .28$, $p = .60$, $\eta p^2 = .02$ | $BF^{01} = 3.1 \pm 1.3\%$<br>$BF_{inclusion} = .12$ | |
| Adjacency x Disc. level | $F_{2,28} = .81$, $p = .43$, $\eta p^2 = .05$ | $BF^{01} = 8.0 \pm 4.8\%$<br>$BF_{inclusion} = .01$ | |
| Adjacency x Disc. sign | $F_{1,14} = .004$, $p = .95$, $\eta p^2 < .001$ | $BF^{01} = 4.4 \pm 6.6\%$<br>$BF_{inclusion} = .004$ | |
| Adj. x Disc. level x Disc. sign | $F_{2,28} = .44$, $p = .65$, $\eta p^2 = .03$ | $BF^{01} = 7.1 \pm 20.1\%$<br>$BF_{inclusion} = .01$ | |
| **Exp. 4** averaging as a function of discrepancy: | | | |
| Discrepancy level (within) | $F_{2,28} = .02$, $p = .97$, $\eta p^2 = .002$ | $BF^{01} = 17.8 \pm 1.0\%$<br>$BF_{inclusion} = .03$ | |
| Discrepancy sign (within) | $F_{1,14} = 2.4$, $p = .14$, $\eta p^2 = .15$ | $BF^{01} = 1.0 \pm 1.1\%$<br>$BF_{inclusion} = .53$ | |
| Homology x Disc. level | $F_{2,28} = .67$, $p = .52$, $\eta p^2 = .05$ | $BF^{01} = 6.5 \pm 2.7\%$<br>$BF_{inclusion} = .02$ | |
| Homology x Disc. sign | $F_{1,14} = .002$, $p = .87$, $\eta p^2 = .001$ | $BF^{01} = 4.4 \pm 9.7\%$<br>$BF_{inclusion} = .28$ | |
| Hom. x Disc. level x Disc. sign | $F_{2,28} = 1.2$, $p = .32$, $\eta p^2 = .08$ | $BF^{01} = 3.8 \pm 3.6\%$<br>$BF_{inclusion} < .001$ | |
| PRECISION | | | |
| **Experiments 1 and 2** single- vs. double- condition: | | | |
| Experiment (Exp.1 vs. Exp.2; between) | $F_{1,28} < .001$, $p = 1.0$, $\eta p^2 < .001$ | $BF^{01} = 2.7 \pm .6\%$<br>$BF_{inclusion} = 1.4$ | |
| Condition (single vs. double; within) | $F_{1,28} = .43$, $p = .51$, $\eta p^2 = .02$ | $BF^{01} = 3.3 \pm 2.2\%$<br>$BF_{inclusion} = 1.3$ | |
| Experiment x Condition | $F_{1,28} = 11.6$, $p = .002$, $\eta p^2 = .29$ | $BF^{01} = .05 \pm 8.4\%$<br>$BF_{inclusion} = 5.0$ | |
| Follow-up (paired t-tests): | Effect of condition in Exp.1: $p = .13$, $d = -.41$;<br>Effect of condition in Exp.2: $p = .001$, $d = 1.1$ | | Greater precision for double-finger condition, but only in bimanual experiment |
| **Experiment 3** single- vs. double-condition: | | | |
| Adjacency (within) | $F_{1,14} = 3.7$, $p = .07$, $\eta p^2 = .21$ | $BF^{01} = 1.2 \pm .9\%$<br>$BF_{inclusion} = .7$ | |
| Condition (single vs. | $F_{1,14} = .73$, $p = .41$, $\eta p^2 = .05$ | $BF^{01} = 2.3 \pm .8\%$ | |

| | | | |
|---|---|---|---|
| double; within) | | $BF_{inclusion}$ = .3 | |
| Adjacency x Condition | $F_{1, 14}$ = .48, $p$ = .50, $\eta p^2$ = .03 | $BF^{01}$ = 2.8 ± 4.9%<br>$BF_{inclusion}$ = .2 | |
| **Experiment 4** single- vs. double-condition: | | | |
| Homology (within) | $F_{1, 14}$ = 1.3, $p$ = .27, $\eta p^2$ = .09 | $BF^{01}$ = 2.4 ± 2.6%<br>$BF_{inclusion}$ = .4 | |
| Condition (single vs. double; within) | $F_{1, 14}$ = 9.0, $p$ = .01, $\eta p^2$ = .39 | $BF^{01}$ = .03 ± 1.8%<br>$BF_{inclusion}$ = 24.4 | |
| Homology x Condition | $F_{1, 14}$ = .002, $p$ = .96, $\eta p^2$ < .001 | $BF^{01}$ = 2.7 ± 2.8%<br>$BF_{inclusion}$ = .4 | |
| **Exp. 1 and 2** averaging as a function of discrepancy: | | | |
| Average direction (within) | $F_{2, 56}$ = .34, $p$ = .71, $\eta p^2$ = .01 | | |
| Experiment x Disc. Level x Disc. Sign x Average | $F_{4, 112}$ = .18, $p$ = .95, $\eta p^2$ = .006 | | |
| Average x Disc. level* | $F_{4, 112}$ = 3.6, $p$ = .008, $\eta p^2$ = .11 | | |
| Follow-up (bonferroni corrected pairwise tests): | Effect of disc. when average was 10: more precise when disc. was 10 than 20 ($p$ = .001) and 30 ($p$ < .001);<br>Effect of disc. when average was 0: all $p$'s = 1.00;<br>Effect of disc. when average -10: all $p$'s = 1.00 | | Precision is higher when discrepancy is low (10), but only when average direction was 10. |
| **Exp. 3** averaging as a function of discrepancy: | | | |
| Average direction (within) | $F_{2, 28}$ = .21, $p$ = .81, $\eta p^2$ = .01 | | |
| Adjacency x Disc. Level x Disc. Sign x Average | $F_{4, 56}$ = 1.6, $p$ = .18, $\eta p^2$ = .10 | | |
| **Exp. 4** averaging as a function of discrepancy: | | | |
| Average direction (within) | $F_{2, 28}$ = 1.1, $p$ = .35, $\eta p^2$ = .07 | | |
| Homology x Disc. Level x Disc. Sign x Average | $F_{4, 56}$ = 1.4, $p$ = .25, $\eta p^2$ = .09 | | |
| Average x Disc. sign | $F_{2, 28}$ = 3.8, $p$ = .03, $\eta p^2$ = .21 | | |
| Follow-up (bonferroni corrected pairwise tests): | Effect of sign when average is 10: $p$ = .56;<br>Effect of sign when average is 0: $p$ = .13;<br>Effect of sign when average is -10: $p$ = .68 | | Effect did not survive the correction for multiple comparisons |

*Note: For the effects of discrepancy, only relevant or statistically significant effects are reported*

**S2: Single-finger conditions.** We conducted preliminary analyses on single-finger conditions separately to establish whether participants were able to perceive component directions and whether perception was similar across individual fingers. Figure S1A shows individual linear regressions fit for each single-finger condition separately in each experiment. The slopes of the regressions represent perceptual sensitivity to component directions and the intercepts of the regressions represents perceptual bias (i.e., the perceived midline of the finger). In addition, standard deviations for repeated judgments of the same component direction represent precision. Table S1 shows the summary of those measures.

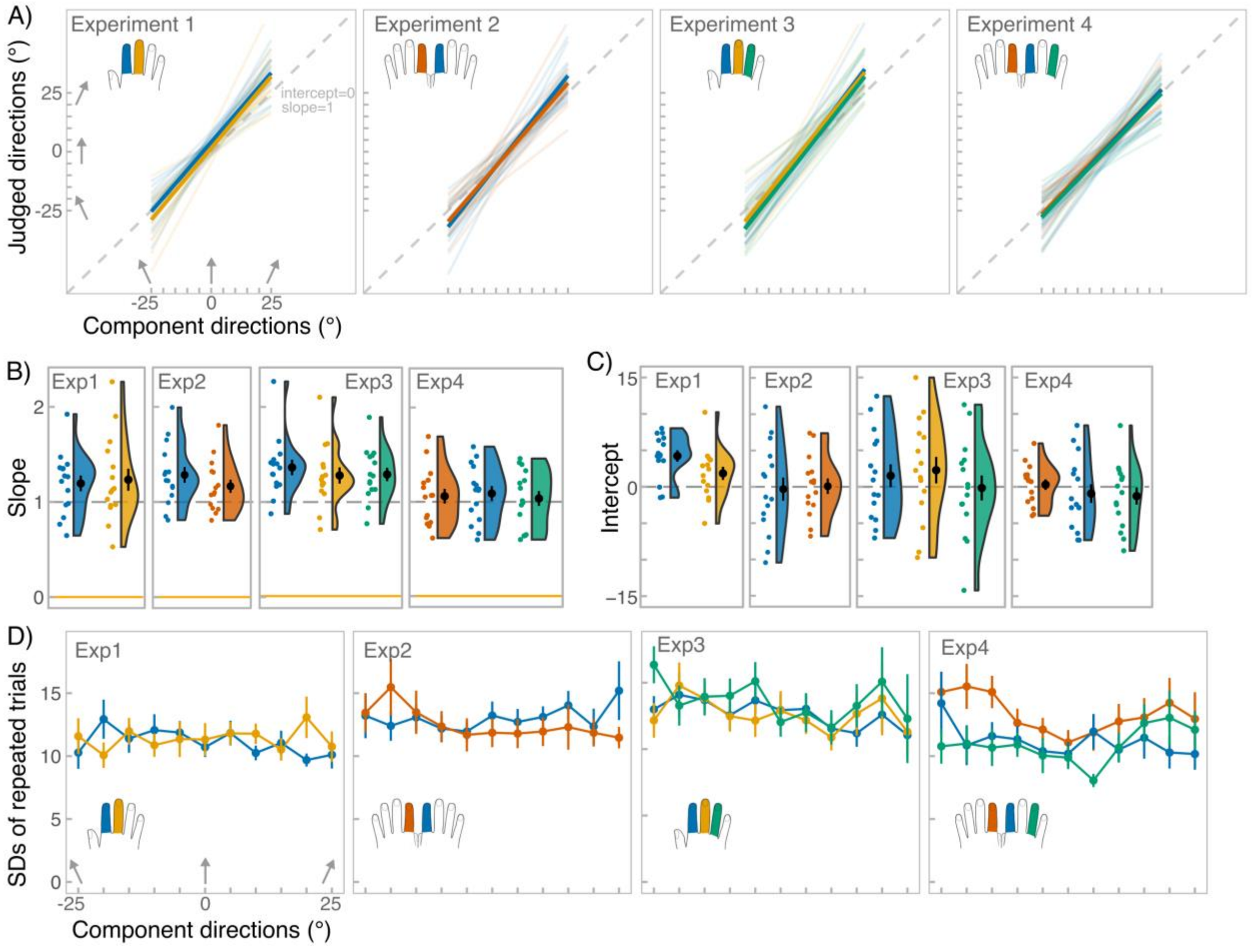


**Figure S2**. Results for single-finger conditions. **A**, shows individual fitted linear regressions (transparent lines) and group-level regressions (thick lines) for each finger, separately for each experiment. The grey dashed line represents equality line of slope 1 and intercept 0. **B**, shows individual slope values (coloured dots) together with the group-level means (black dot). Raincloud plots represent data distribution and error bars represent standard error from the mean. Colours on the plots correspond to the finger-conditions in A. **C**, shows individual intercept values together with the group-level means. **D**, shows group-level standard deviations (SDs) for repeated trials for each component direction. Error bars represent standard error from the mean.

**Table S2:**
Main measures for single-finger conditions

| | Slope | | | Intercept | | SD |
|---|---|---|---|---|---|---|
| Condition | Mean ± SD | t-test against 0: *t*(*d*) | t-test against 1: *t*(*d*) | Mean ± SD | t-test against 0: *t*(*d*) | Mean ± SD |
| **Exp. 1:** | | | | | | |
| Index | 1.2 ± .3 | 14.39(3.72)** | 2.35(.61)* | 4.22 ± 3.1 | 5.35(1.38)** | 11.1 ± 4.1 |
| Middle | 1.2 ± .5 | 10.66(2.75)** | 2.02(.52) | 1.84 ± 3.5 | 2.04(.53) | 11.4 ± 4.5 |
| **Exp. 2:** | | | | | | |
| Left index | 1.3 ± .3 | 16.70(4.31)** | 2.37(61)* | 0.09 ± 4.1 | .08(.02) | 13.0 ± 5.1 |
| Right index | 1.2 ± .3 | 15.43(3.98)** | 3.42(.88)* | -0.3 ± 6.2 | -.20(-.05) | 12.5 ± 5.5 |
| **Exp. 3:** | | | | | | |
| Index | 1.4 ± .3 | $120^{\dagger}$(4.32)** | $118^{\dagger}$(1.15)** | 1.5 ± 6.1 | .97(.25) | 12.7 ± 4.5 |
| Middle | 1.3 ± .3 | 15.02(3.88)** | 3.28(0.85)* | 2.3 ± 7.0 | 1.26(.33) | 12.6 ± 6.1 |
| Ring | 1.3 ± .3 | 16.26(4.20)** | 4.11(1.06)* | -0.2 ± 6.6 | -.09(-.02) | 13.6 ± 6.9 |
| **Exp. 4:** | | | | | | |
| Left index | 1.1 ± .3 | 13.49(3.5)** | .75(.19) | 0.3 ± 2.8 | .41(.11) | 13.3 ± 5.9 |
| Right index | 1.1 ± .3 | 13.91(3.6)** | 1.12(.29) | -0.9 ± 5.0 | -.72(-.19) | 11.2 ± 5.3 |
| Right ring | 1.0 ± .3 | 13.16(3.4)** | .45(.12) | -1.3 ± 4.5 | -1.09(-.28) | 10.9 ± 5.3 |

*p < .05, **p < .001. $x^{\dagger}$ indicate the sign test (V), which was used instead of t-test, due to non-normality. Same group of 15 participants performed under all single-finger conditions within a single experiment, but each experiment had a new group of 15 participants.

Figure S2B shows single-subject and group-level slope values for each finger across all four experiments. The slopes were significantly above 0 in all experiments indicating that the component directions could be successfully perceived. In fact, in all experiments slopes were either not significantly different from 1, indicating perfect sensitivity, or greater than 1, showing overestimation. We compared sensitivity between fingers. In Experiment 1, sensitivity did not differ between index and middle fingers (paired-sample t-test: $t_{14}$ = -0.64, $p$ = .53, $d$ = -.17). Similarly, in Experiment 2 sensitivity was not significantly different between right and left index fingers ($t_{14}$ = 1.89 $p$ = .08, $d$ = .49). In Experiment 3, one-way repeated-measures ANOVA yielded a significant effect of finger ($F_{2, 28}$ = 3.33, $p$ = .05, $\eta_p^2$ = .19). We expected perception to be possibly worse on the ring finger compared to index and middle fingers. However, planned contrasts showed that ring finger did not differ from index and middle fingers ($t_{28}$ = 1.03, $p$ = .31). Rather the slope for index finger judgments was higher than for middle finger judgments ($t_{28}$ = 2.37, $p$ = .03). Yet, the effect was marginal and all fingers yielded slope values greater than 1. Lastly, in Experiment 4, there was no significant effect of finger ($F_{2, 28}$ = .54, $p$ = .59, $\eta_p^2$ = .14).

Figure S2C shows single-subject and group-level intercept values for each finger, separately for each experiment. Except for the intercept for index finger in Experiment 1, all other intercepts did not differ from 0 significantly, indicating no bias. Judgments on index finger in Exp. 1 showed a positive bias of ~ 4°, meaning that the perceived midline of the finger was shifted slightly rightwards. Indeed, there was a significant difference between single-finger intercepts in Exp. 1 (sign test due to non-normality: *V= 99*, $p$ = .03, $d$ = 0.66). There was no between finger difference in Exp. 2 (paired-sample t-test: $t_{14}$ = -.19, $p$ = .85, $d$ = -.05). Similarly, there was no effect of finger in Exp. 3 ($F_{2, 28}$ = 1.50, $p$ = .24, $\eta_p^2$ = .10) or Exp. 4 ($F_{2, 28}$ = .71, $p$ = .50, $\eta_p^2$ = .05).

For precision, we calculated unbiased SDs for each component direction separately for each finger (Fig. S2D) and ran a rmANOVA with factors finger (stimulated fingers in the particular experiment), overall direction (leftward vs. rightward) and specific angle (25°, 20°, 15°, 10°, 5°). We excluded the component direction 0° from this analysis, as it could not be divided into leftward vs. rightward direction. In Experiment 1, there was no significant effect of finger ($F_{1, 14}$ = .18, $p$ = .68, $\eta_p^2$ = .01), indicating that both fingers were equally reliable. The effects of overall direction and specific direction were also non-significant ($F_{1, 14}$ = .31, $p$ = .58, $\eta_p^2$ = .02 and $F_{4, 56}$ = .43, $p$ = .78, $\eta_p^2$ = .02, respectively). All interactions were non-significant as well (p values: .16 > p > .90). No effect of finger ($F_{1, 14}$ = .27, $p$ = .61, $\eta_p^2$ = .02) was revealed in Experiment 2. Other main effects and interactions also remained non-significant (all other p values: .08 > p > .79). In Experiment 3, effect of finger was non-significant ($F_{2, 28}$ = 1.53, $p$ = .23, $\eta_p^2$ = .10), but there was a significant effect of overall direction ($F_{1, 14}$ = 9.29, $p$ = .009, $\eta_p^2$ = .40) with participants being more precise judging leftward (mean SD= 12) relative to rightward (mean SD = 14) directions. Lastly, in Experiment 4, we did observe a significant effect of finger ($F_{2, 28}$ = 6.1, $p$ = .007, $\eta_p^2$ = .30) with judgments from left index finger being more variable (mean SD = 13) than both right index (mean SD = 11) and right ring (mean SD = 11). In addition, the effect of specific angle was also significant ($F_{4, 56}$ = 3.1, $p$ = .015, $\eta_p^2$ = .18 - Greenhouse-corrected p-value), Bonferroni-corrected pairwise tests indicating a significant difference only between 5° and 15° as well as 5° and 25° with judgements about 5° being more precise (mean SD = 11) than for 15° (mean SD = 12) and 25° (mean SD= 12). Note that 0° was excluded from analysis, but its mean SD was also low (mean SD = 10). This suggests that precision might have been higher, when estimating directions that were around the midline of the finger relative to the ones that deviated away from the midline. But this effect was only present in Experiment 4.

In summary, participants can successfully extract component directions in all experiments and their judgments seem to be generally unbiased (except in one case; Exp.1 index finger). Within experiments, considering all measures (slope, intercept, SD), perception across different fingers was similar with few exceptions: increased bias on index in Exp.1 and reduced precision on left finger in Exp.4. Analysis of SDs showed that there were a few direction-specific effects, namely in Exp.3 and Exp.4, however overall the direction-related differences were small.

**S3: Full statistical analysis of weighting data derived from data and model**

**Table S3**
Full statistical analysis of weighting data derived from data and model

| | | |
|---|---|---|
| WEIGHTING DERIVED FROM THE DATA | | |
| **Unimanual (Adj; Exp1 & Exp3)** | | |
| Average (within) | $F_{2, 56}$ = 10.0, $p$ = .001, $\eta p^2$ = .26 | |
| Follow-up (bonferroni paired tests) | Weight higher when average is 10 compared to when it's -10 ($p$ = .004) | Virtually leading finger is weighted higher in average direction estimation |
| Experiment x Average | $F_{2, 56}$ = .27, $p$ = .77, $\eta p^2$ = .01 | |
| Exp. x Av. x Disc. level x Disc. sign | $F_{4, 112}$ = .86, $p$ = .49, $\eta p^2$ = .03 | |
| Disc. level x Disc. Sign* | $F_{2, 56}$ = 15.7, $p$ = .002, $\eta p^2$ = .36 | |
| Follow-up (bonferroni paired tests) | effect of disc. level when discrepancy is positive: weight lower when disc. is 10 relative to 20 ($p$ < .001) and 30 ($p$ < .001)<br>effect of disc. level when discrepancy is negative: weight higher when disc. is 10 relative to 20 ($p$ < .001) and 30 ($p$ < .001) | Discrepancy between component directions modulates weighting, but in opposite directions depending on whether components are converging or diverging. |
| **Unimanual (Non-adj; Exp3)** | | |
| Average (within) | $F_{2, 28}$ = 6.7, $p$ = .006, $\eta p^2$ = .32 | |
| Follow-up (bonferroni paired tests) | Weight higher when average is 10 compared to when it's -10 ($p$ = .001) | Virtually leading finger is weighted higher in average direction estimation |
| Av. x Disc. level x Disc. sign | $F_{4, 56}$ = .23, $p$ = .92, $\eta p^2$ = .02 | |
| **Bimanual (Hom; Exp2 & Exp4)** | | |
| Average (within) | $F_{2, 56}$ = .43, $p$ = .62, $\eta p^2$ = .02 | |
| Experiment x Average | $F_{2, 56}$ = .36, $p$ = .67, $\eta p^2$ = .01 | |
| Exp. x Av. x Disc. level x Disc. sign | $F_{1, 28}$ = 1.30, $p$ = .28, $\eta p^2$ = .04 | |
| Average x Disc. Sign* | $F_{2, 56}$ = 3.9, $p$ = .04, $\eta p^2$ = .12 | |
| Follow-up (bonferroni paired tests) | Weighting modulated with average directions, but only when directions were diverging ($p$ = .01) | |
| **Bimanual (Non-hom; Exp4)** | | |
| Average (within) | $F_{2, 28}$ = .33, $p$ = .72, $\eta p^2$ = .02 | |
| Av. x Disc. level x Disc. sign | $F_{4, 56}$ = .66, $p$ = .64, $\eta p^2$ = .04 | |
| WEIGHTING DERIVED FROM THE MODEL | | |
| **Unimanual (Adj; Exp1 & Exp3)** | | |
| Experiment (between) | $F_{1\ 28}$ = 2.1, $p$ = .16, $\eta p^2$ = .07 | |

| | | |
|---|---|---|
| Average (within) | $F_{2, 56} = 20.8$, $p < .001$, $\eta p^2 = .43$ | |
| Follow-up (bonferroni paired tests) | Weight higher when average is -10 compared to when it's 10 ($p$ < .001 | Virtually leading finger is weighted higher in average direction estimation |
| Discrepancy (within) | $F_{6, 168} = .84$, $p = .54$, $\eta p^2 = .03$ | |
| Average x Discrepancy | $F_{12, 336} = 6.2$, $p < .001$, $\eta p^2 = .18$ | |
| Follow-up (ANOVAs) | Effect of disc. when average is -10: $p = .74$, $\eta p^2 = .02$; Effect of disc. when average is 0: $p = .47$, $\eta p^2 = .03$; Effect of disc. when average is 10: $p = .001$, $\eta p^2 = .12$ | Discrepancy modulates weighting, but only when average is 10, however, effect is small. |
| **Unimanual (Non-adj; Exp3)** | | |
| Average (within) | $F_{2, 28} = 18.94$, $p < .001$, $\eta p^2 = .58$ | |
| Follow-up (bonferroni paired tests) | Weight higher when average is -10 compared to when it's 10 ($p$ < .001) | Virtually leading finger is weighted higher in average direction estimation |
| Discrepancy (within) | $F_{6, 84} = .15$, $p = .99$, $\eta p^2 = .01$ | |
| Average x Discrepancy | $F_{12, 168} = 7.5$, $p < .001$, $\eta p^2 = .35$ | |
| Follow-up (ANOVAs) | Effect of disc. when average is -10: $p = .03$, $\eta p^2 = .15$; Effect of disc. when average is 0: $p = 1.0$, $\eta p^2 = .001$; Effect of disc. when average is 10: $p = .67$, $\eta p^2 = .05$ | Effect does not survive multiple comparisons. |
| **Bimanual (Hom; Exp2 & Exp4)** | | |
| Average (within) | $F_{2, 56} = 1.6$, $p = .21$, $\eta p^2 = .05$ | |
| Discrepancy (within) | $F_{6, 168} = 2.2$, $p = .05$, $\eta p^2 = .07$ | |
| Average x Discrepancy | $F_{12, 336} = 18.7$, $p < .001$, $\eta p^2 = .40$ | |
| Follow-up (ANOVAs) | Effect of disc. when average is -10: $p < .001$, $\eta p^2 = .21$; Effect of disc. when average is 0: $p = .04$, $\eta p^2 = .08$; Effect of disc. when average is 10: $p = 1.0$, $\eta p^2 < .001$ | Discrepancy modulates weighting, but only when average is -10. |
| **Bimanual (Non-hom; Exp4)** | | |
| Average (within) | $F_{2, 28} = 5.0$, $p = .02$, $\eta p^2 = .26$ | Effect does not survive multiple comparisons. |
| Discrepancy (within) | $F_{6, 84} = .86$, $p = .53$, $\eta p^2 = .06$ | |
| Average x Discrepancy | $F_{12, 168} = .92$, $p = .53$, $\eta p^2 = .06$ | |

**S4: Effect of judgement biases on the estimation of weights.** When the estimate of motion angle for each finger is distorted as shown in the following equation

$$\tilde{\theta}_i = \theta_i + b \quad \text{(S4.1)}$$

the estimated weight of the left-most finger ($\widehat{w}_1$) would be given using the actual weight given to the left-most finger ($w_1$) as follows

$$\widehat{w}_1 = \frac{\widehat{\theta}_{avr} - \theta_2}{\theta_1 - \theta_2} = w_1 - \frac{\mathrm{b}}{\theta_{dsc}} \quad \text{(S4.2)}$$

This shows that the bias in the judgements can cause apparent effects of discrepancy on the estimated weights.

Note that the equation explains the effect of discrepancy on the apparent weight found for the adjacent condition. Namely, when the component directions are diverging ($\theta_{dsc} > 0$), the weight would be lower when discrepancy is low (10°) compared to when it was high (20° and 30°). Opposite pattern is expected when component directions are converging ($\theta_{dsc} < 0$); i.e., weight would be higher when discrepancy is low compared to when it is high.

**S5: Condition for prioritising the leading finger in the biased integration model.** The condition for the biased integration model to assign more weight to the virtual leading finger (defined by average angle) can be given as follows.

$$\begin{cases} w_1 > \frac{1}{2}\ (if\ \theta_{avr} > 0) \\ w_1 < \frac{1}{2}\ \ (if\ \theta_{avr} < 0) \end{cases} \tag{S5.1}$$

Using equations (9) and (10), this is equivalent to

$$\begin{cases} c_1\theta_1 > c_2\theta_2\ (if\ \theta_{avr} > 0) \\ c_1\theta_1 < c_2\theta_2\ \ (if\ \theta_{avr} < 0) \end{cases} \tag{S5.2}$$

Using the following definitions also found in the paper

$$\theta_{dsc} = \theta_2 - \theta_1 \tag{S5.3}$$

$$\theta_{avr} = (\theta_1 + \theta_2)/2 \tag{S5.4}$$

$$\bar{c} = (c_1 + c_2)/2 \tag{S5.5}$$

$$\Delta c = (c_1 - c_2) \tag{S5.6}$$

the condition can be transformed as follows.

$$\begin{cases} \Delta c\theta_{avr} > \bar{c}\theta_{dsc}\ (if\ \theta_{avr} > 0) \\ \Delta c\theta_{avr} < \bar{c}\theta_{dsc}\ \ (if\ \theta_{avr} < 0) \end{cases} \tag{S5.7}$$

The condition for the model to satisfy the above conditions for all discrepancies is as follows.

$$\bar{c} = 0 \tag{S5.8}$$

$$\Delta c > 0 \tag{S5.9}$$

Namely, the two fingers should have gain factors with an average of zero and a positive difference in order to assign more weight to the leading finger.

**S6: Judgement prediction of the biased-integration model.** Following figures show the judgement prediction of the biased-integration model.

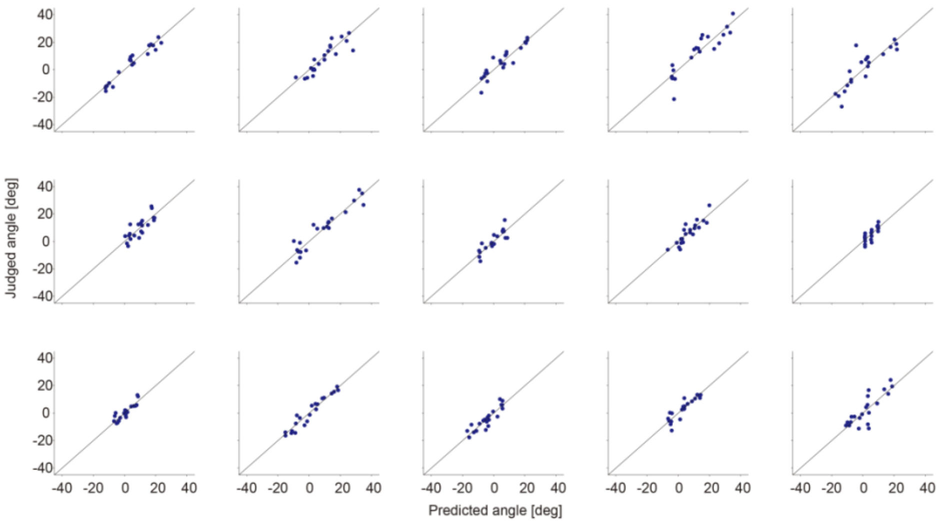


**Figure S6.1** Judgement prediction for Experiment 1.

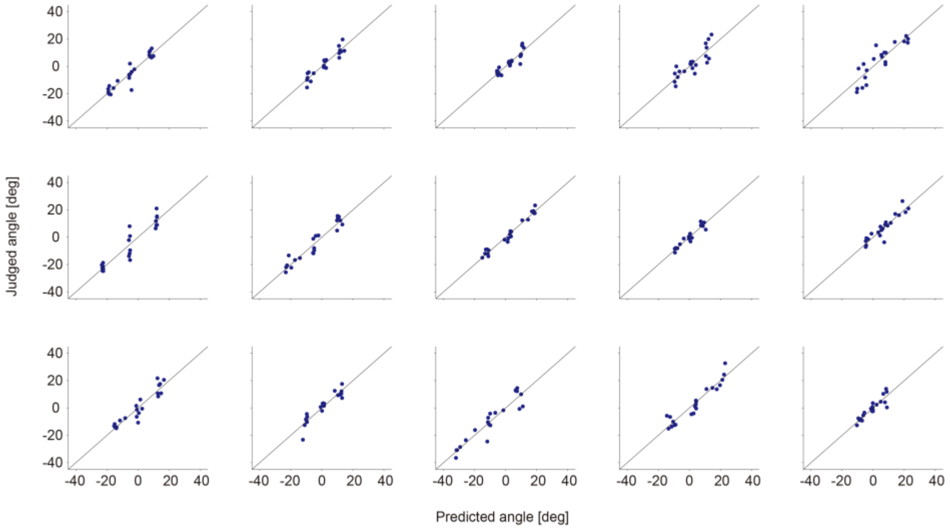


**Figure S6.2** Judgement prediction for Experiment 2.

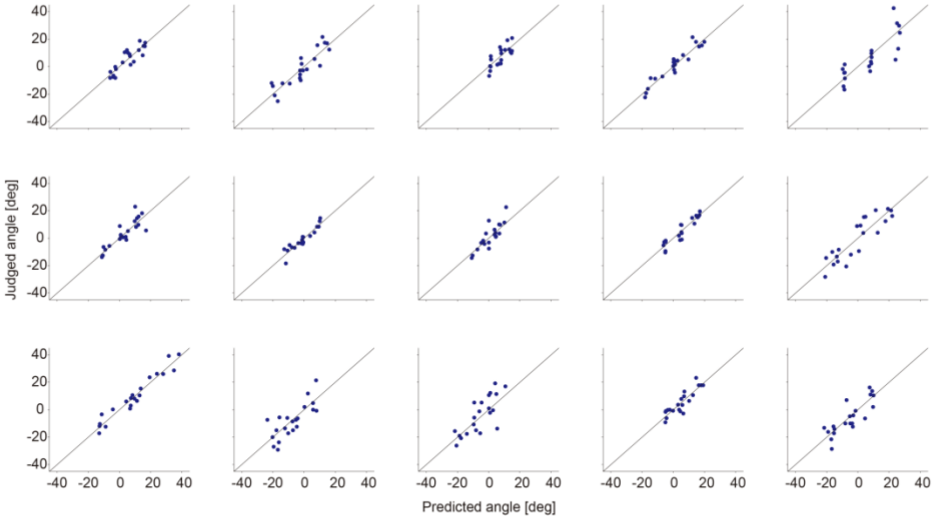


**Figure S6.3** Judgement prediction for Experiment 3 (Adjacent).

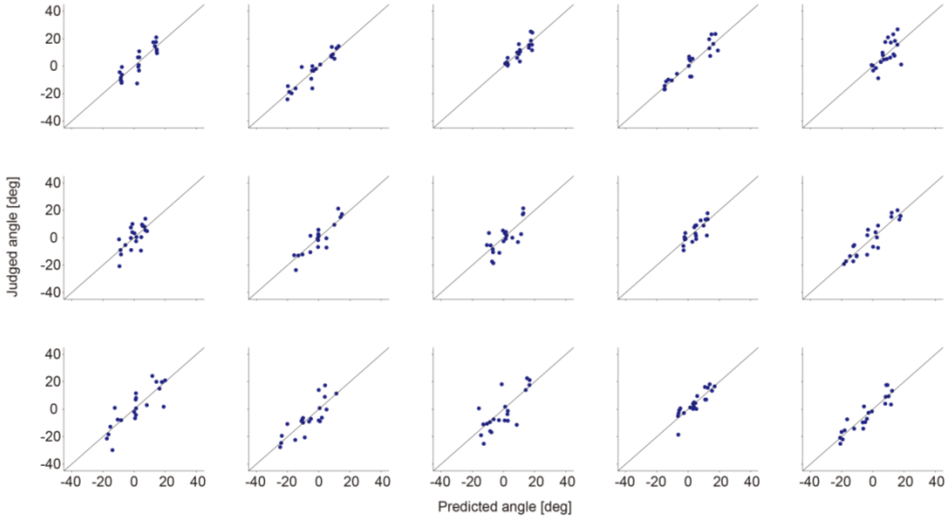


**Figure S6.4** Judgement prediction for Experiment 3 (Non-adjacent).

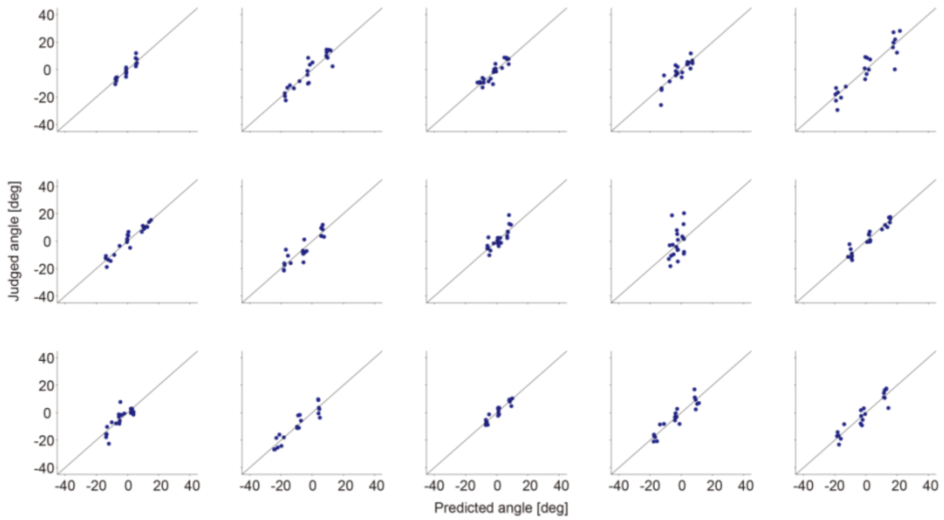


**Figure S6.5** Judgement prediction for Experiment 4 (Homologous)

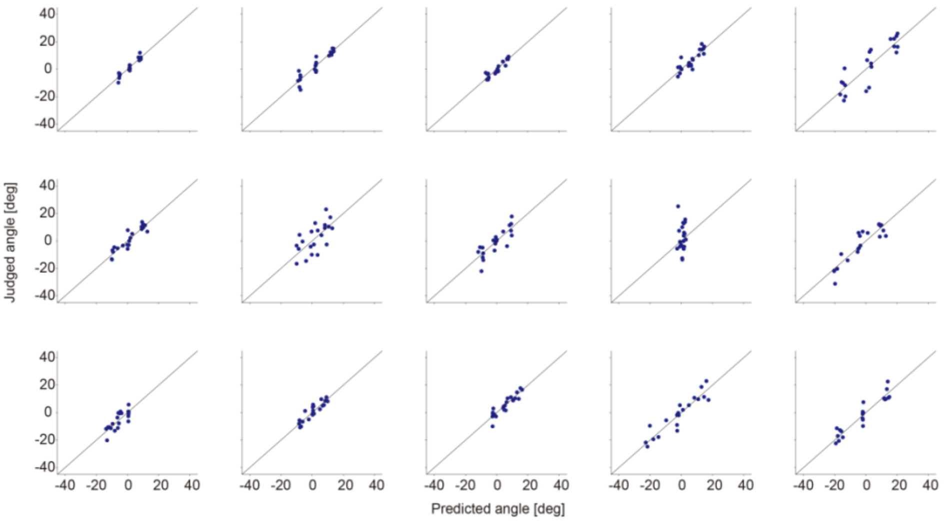


**Figure S6.6** Judgement prediction for Experiment 4 (Non-homologous)

**S7: Weight assignment of the model**

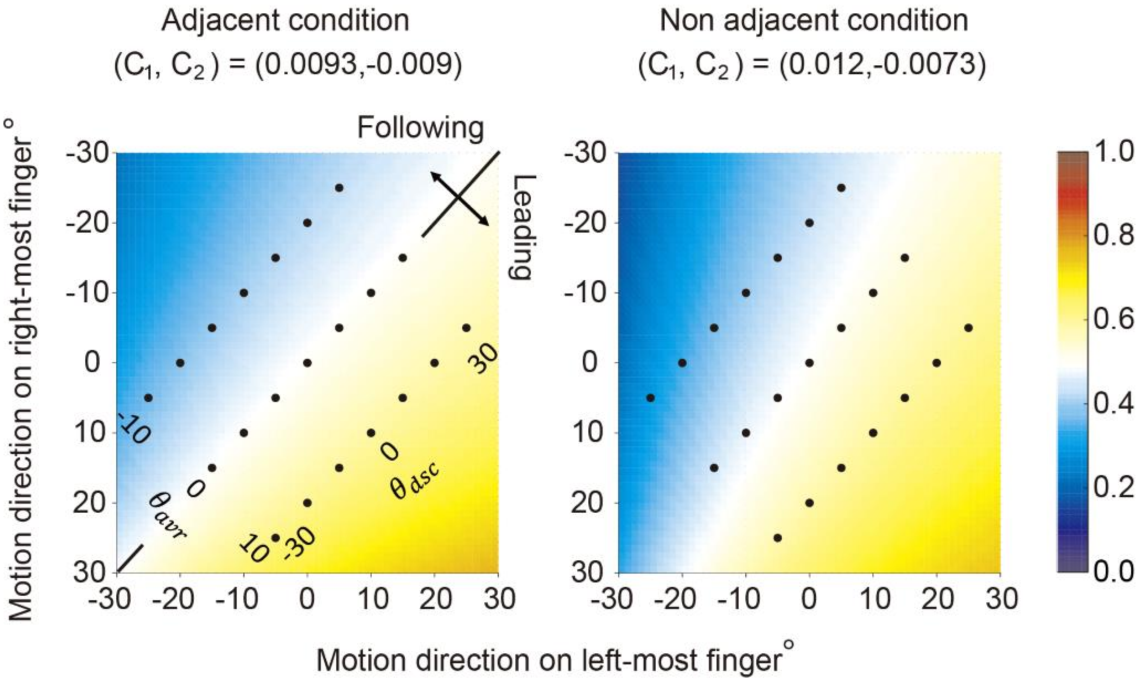


**Figure S7**. Weight assignment calculated from the estimated gain factors averaged across participants (left panel: adjacent condition, right panel: non-adjacent condition). The colors in the figure show the weight that the model assigns to the left-most finger for different set of gain factors. Black dots show the stimulus pairs used in the experiment. Dots aligned from bottom left to top right have same average angles, whereas dots aligned from bottom right to top left have same discrepancies of angles. X and Y axes are aligned to Figure 1C (see manuscript).

When gain factors were tuned to predict the data in unimanual experiments, adjacent fingers had similar impact on the model weight assignment (paired t-test between c1 and c2: $t_{29} = 0.1$, p = .92, d =0.02; mean c1 = .0093 ± .018 & mean c2 = -.0090 ± .017; Fig. S7 left). In non-adjacent pairs, index finger seemed to have a slightly larger impact on weight assignment, but the difference was not statistically significant (paired t-test between c1 and c2: $t_{14} = 0.79$, p = .45, d =0.2; mean c1 = .012 ± .017 & mean c2 = -.0073 ± .014; Fig. S7 right).

As with the weights derived from the data, we examined whether model weights also varied as a function of average angle (for breakdown of all effects see Table S3). Because the model could estimate the weights when discrepancy was 0, we did not add sign of discrepancy as a separate effect, but only evaluated the effect of level of discrepancy (-30°, -20°, -10°, 0°, 10°, 20°, 30°). Both when unimanual finger were adjacent and non-adjacent, analysis revealed a significant effect of average angle ($F_{2, 56}$ = 20.85, p < .001 $\eta p^2$ = .43 & $F_{2, 28}$ = 18.94, p = .001, $\eta p^2$ = .58, respectively). Post-hoc Bonferroni-corrected pairwise tests showed that the model weight was significantly larger for stimuli with average angle of 10° compared to those with average angle of -10° (p < .001 for both adjacent and non-adjacent fingers). Model weights did not vary with discrepancy (main effect of discrepancy: $F_{6, 168}$ = .84, p = .54, $\eta p^2$ = .01 & $F_{6, 84}$ = .15, p = .99, $\eta p^2$ = .01, respectively for adjacent and non-adjacent fingers). This implies that, when considering the biases in the individual estimates, the fitted model assigned more weight to the virtual leading finger. In contrast, for bimanual experiments the model weights were not modulated by average angle, when fingers were homologous ($F_{2, 56}$ = 1.59, p = .21, $\eta p^2$ = .05). When fingers were non-homologous, the effect of angle reached significance ($F_{2, 28}$ = 5.03, p = .02, $\eta p^2$ = .26), however considering multiple ANOVA's, it would not survive Bonferroni correction.

**S8: Relation between biased sensory weighting and decrease in precision.** We found that sensory weighting among the two fingers are biased in the unimanual conditions. Since such biases in sensory weighting could decrease the precision in the averaging task, we examined whether the estimated biases account for the unimanual-bimanual difference in the precision found in our study.

**Bias in sensory weighting can decrease precision in averaging task.** First, we explain why the bias in sensory weighting can result in decreased precision. In all our experiments, the two fingers had similar precision (i.e., similar SD in judgements). Furthermore, analysis of SDs across component directions did not show any consistent direction-specific effects. We can therefore assume that each finger (i=1,2) produced an estimate ($\hat{\theta}_i$) of the component direction ($\theta_i$) with similar variance ($\sigma^2$) across all component directions. Considering the distribution of the judgements, the distribution of the estimates can be modeled as a Gaussian distribution.

$$p\left(\hat{\theta}_i\right) = N(a_i\theta_i + b_i, \sigma^2) = \frac{1}{\sqrt{2\pi}\sigma} exp\left(-\frac{(a_i\theta_i+b_i-\hat{\theta}_i)^2}{2\sigma^2}\right) \quad \text{(S8.1)}$$

Note that we also consider the distortion of the estimate both in terms of sensitivity ($a_i$) and bias ($b_i$). In our study, participants were instructed to average the directions of the motion stimuli delivered to the two fingers. Therefore, we can assume that the judgements were based on weighted average of the two estimates as follows.

$$\theta_{jud} = \sum_{i=1}^{2} w_i\tilde{\theta}_i \quad \text{(S8.2)}$$

$$\sum_{i=1}^{2} w_i = 1 \quad \text{(S8.3)}$$

When the variance of the two estimates are independent, SD of the judgment can be predicted as follows.

$$\hat{\sigma}_{jud} = \mathrm{K}(w_i)\sigma \quad \text{(S8.4)}$$

$$\mathrm{K}(w_i) = \sqrt{\sum_{i=1}^{2} w_i^2} \quad \text{(S8.5)}$$

$\mathrm{K}(w_i)$ represents the gain in SD caused by integrating the signals. Note that the distortion in the judgements ($a_i, b_i$) are no longer relevant. Fig. S8.1 plots $\mathrm{K}(w_i)$ against the weight given to the first finger. This shows that SD of the judgment would be minimum ($1/\sqrt{2}$) when the weights are equal ($w_1 = w_2 = 1/2$), but will increase as the bias in the weights ($w_{bias} = |w_1 - 0.5|$) increases.

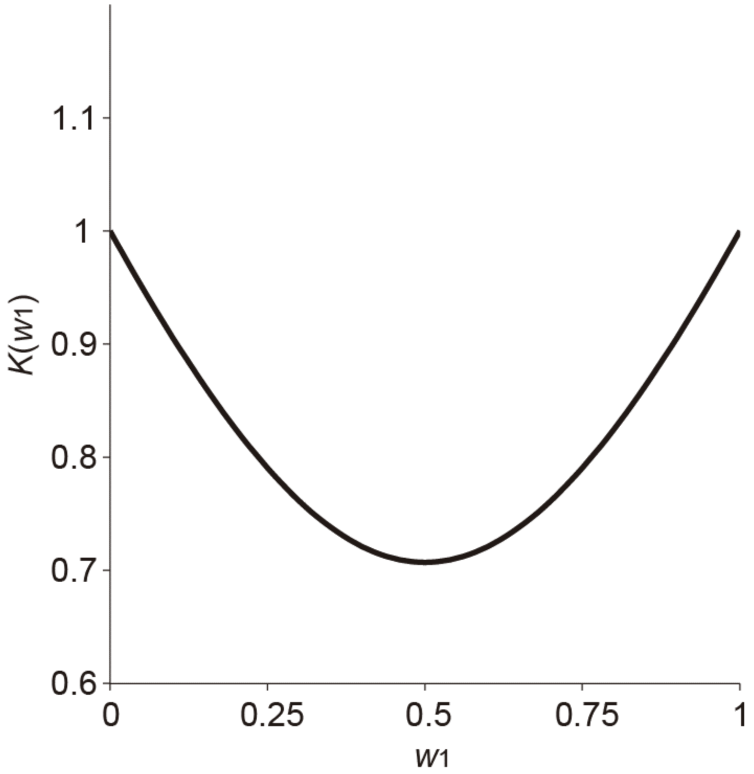


**Fig. S8.1** Relation between SD and weight

**Bias in estimated weight do not account for unimanual-bimanual difference in precision.** Next, we show that the estimated biases in the sensory weighting do not account for the difference in precision between the unimanual and bimanual conditions. To this end, we first estimated the weight given to the the left-most finger (right index finger in unimanual conditions and left index finger in bimanual conditions) by fitting our computational model (biased integration model; see companion paper for detail) to the judgements. Then, we calculated the gain in SD based on equation (S8.5). Table S8.1 summarizes the gain averaged across all participants and stimulus pairs for each experiment.

**Table S8.1**

Summary of estimated gain in SD due to bias in weight

| | **Exp. 1** | **Exp. 3** | **Exp. 2** | **Exp. 4** |
|---|---|---|---|---|
| Each Exp. | 0.76 (0.01) | 0.75 (0.01) | 0.72 (0.01) | 0.72 (0.01) |
| Avr. Across Exp. | 0.75 (0.01) | | 0.72 (0.01) | |

*Note: values represent mean (SE)*

**Table S8.2**

Summary of gain in SD measured in the experiments

| | Exp. 1 | Exp. 3 | Exp. 2 | Exp. 4 |
|---|---|---|---|---|
| Each Exp. | 1.16 (0.07) | 1.00 (0.06) | 0.91 (0.04) | 0.90 (0.04) |
| Avr. Across Exp. | 1.08 (0.05) | | 0.91 (0.03) | |

*Note: values represent mean (SE)*

To match the summary data of SD shown in Figure 7, we aggregated data between adjacency and homology in experiments 3 and 4. Bottom row shows the gain averaged across the two experiments. The ratio of the averaged gain factor between the unimanual and bimanual condition (.75/.72) predicts that the SD in the unimanual condition would be approximately 4% larger than the bimanual condition. In contrast to this prediction, the SD in the unimanual condition was approximately 17% larger than that of the bimanual condition (Fig. 7). Furthermore, the ratio of SD between double finger condition and single finger condition ($\sigma_{jud}/\ \sigma$ ; Table S8.2), which corresponded to the actual gain $\mathrm{K}(w_i)$, was much larger than that estimated from the weights. This suggests a possibility that some noise was shared among the two fingers. In such case, the ratio of SD between the unimanual and bimanual condition is predicted to be even smaller than 4%. Taken together, the analysis shows that the difference in precision between the unimanual and bimanual conditions cannot be explained based only on the bias in the sensory weighting.

**S9: Difficulty to explain the effect of average angle on weights based on sensitivities and biases**. Let's assume that the two fingers have different sensitivities and biases. The estimate of motion angle of the two fingers can be given as follows.

$$\tilde{\theta}_1 = a_1\theta_1 + b_1 \quad \text{(S9.1)}$$

$$\tilde{\theta}_2 = a_2\theta_2 + b_2 \quad \text{(S9.2)}$$

When the two fingers are equally weighted, the judged average angle can be given using equation (8) as follows.

$$\hat{\theta}_{avr} = \frac{\tilde{\theta}_i+\tilde{\theta}_2}{2} = \frac{a_1\theta_1+a_2\theta_2}{2} + \frac{b_1+b_2}{2} \quad \text{(S9.3)}$$

Therefore, the apparent weight of the left-most finger can be given using equation (1) as follows.

$$\mathrm{w} = \frac{\hat{\theta}_{avr}-\theta_2}{\theta_1-\theta_2} = \frac{a_1\theta_1+(a_2-2)\theta_2}{2(\theta_1-\theta_2)} + \frac{b_1+b_2}{2(\theta_1-\theta_2)} \quad \text{(S9.4)}$$

Using equations (4) and (5), the weight can be represented using the average angle and the discrepancy as follows.

$$\mathrm{w} = \frac{\hat{\theta}_{avr}-\theta_2}{\theta_1-\theta_2} = -\frac{a_1\theta_{avr}+(a_2-a_1-2)(\theta_{avr}+\theta_{dsc}/2)}{2\theta_{dsc}} - \frac{b_1+b_2}{2\theta_{dsc}} \quad \text{(S9.5)}$$

When we average the weight between two conditions with the same average angle and opposite discrepancies, the weight would be as follows.

$$\mathrm{w} = \frac{\hat{\theta}_{avr}-\theta_2}{\theta_1-\theta_2} = \frac{2-a_2+a_1}{4} \quad \text{(S9.6)}$$

The weight does not depend on the average angle. This shows that the effect of average angle on the apparent weight of the left-most finger (Fig. 8A and 8B) cannot be explained by the difference in sensitivity and/or biases.

**S10: Alternative models for the virtual-leading-finger priority (VLF-priority).** Details of the alternative models are described below and AIC values for all the models are summarized in table S10.

**(a) Biased integration model with sensitivity/slope identified in single finger experiment (a and b from 1 finger exp.)** This model was equal to the biased integration model except that it used sensitivity (a) and bias (b) values that was identified for each finger by fitting the judgements made in single finger experiments using the following equation.

$$\theta_{jud} = \hat{a}_i\theta_i + \hat{b}_i \tag{S10.1}$$

The estimated values were then used in the biased integration model to calculate the individual estimates using the following equation.

$$\tilde{\theta}_i = \hat{a}_i\theta_i + \hat{b}_i \tag{S10.2}$$

We found that AIC values were lower for the biased integration model which assumed identical sensitivity and bias for the two fingers.

**(b) Biased integration model with ideal sensitivity (a = 1).** This model was equal to the biased integration model except that it assumed an ideal sensitivity of a=1. The model had larger AIC values than the biased integration model. Therefore, the use of the sensitivity factor (a) was justified.

**(c) Biased integration model with no bias (b = 0).** This model was equal to the biased integration model except that it assumed no bias in estimates of individual fingers (i.e., b=0). The model had larger AIC values than the biased integration model. Therefore, the use of the bias factor (b) was justified.

**(d) Vector summation model.** This model calculated the weights in the same way that the biased-integration model did, but determined the final estimate by vector summation. Namely, vector of tactile motion for each finger was scaled by the allocated weights and the two scaled vectors were combined by vector summation. The final estimate was calculated as the direction of the combined vector. The final estimate was calculated by the following equation.

$$\hat{\theta}_{avr} = atan\left(\frac{\sum_{i=1}^{2} w_i sin\tilde{\theta}_i}{\sum_{i=1}^{2} w_i cos\tilde{\theta}_i}\right) \tag{S10.3}$$

As expected from the fact that weighted averaging of angles and vectors are similar in our case, the AIC value was similar to that of the biased integration model.

**(e) Vector summation model without normalization.** This was similar to the vector summation model except that it skipped the normalization step for calculating the weights. The final estimate was calculated as follows.

$$\hat{\theta}_{avr} = atan(\frac{\sum_{i=1}^{2} g_i sin\tilde{\theta}_i}{\sum_{i=1}^{2} g_i cos\tilde{\theta}_i}) \quad \text{(S10.4)}$$

This model had larger AIC values than the vector summation model and suggested the relevance of the normalization process.

**(f) Lateral interaction model.** This model was similar to the biased integration model. The only difference was that the angle of tactile motion on the *other* finger regulated the strength in which each finger attracted the weight.

$$g_1 = \frac{1}{2} + c_1\tilde{\theta}_2 \quad \text{(S10.5)}$$

$$g_2 = \frac{1}{2} + c_2\tilde{\theta}_1 \quad \text{(S10.6)}$$

The model would allocate more weight to the leading finger when the gain factors $c_1$ is negative and/or $c_2$ is positive. AIC scores were similar to that of the biased-integration model in most cases. However, in the adjacent condition of experiment 2, the biased integration model provided a lower AIC value.

**(g) Lateral inhibition model.** This model was similar to the lateral interaction model. However, while the lateral interaction model allowed both lateral inhibition and excitation, this model was designed to only allow lateral inhibition. The weights were calculated using the following equation.

$$g_1 = \frac{1}{2} - c_1 \max(-\tilde{\theta}_2, 0) \quad \text{(S10.7)}$$

$$g_2 = \frac{1}{2} - c_2 \max(\tilde{\theta}_1, 0) \quad \text{(S10.8)}$$

AIC scores were similar to that of the biased-integration model in most cases. However, in experiment 1, the biased integration model provided a lower AIC value.

**Table S10**
AIC values for all tested models.

| Experiment | Exp. 1 | Exp. 3 | | Exp. 2 | Exp. 4 | |
|---|---|---|---|---|---|---|
| U/B | | Unimanual | | | Bimanual | |
| Condition | Adj. | Adj. | N-adj. | Hom. | Hom. | N-hom. |
| Unbiased integration | **1332.5** | **828.5** | **845.8** | 1292.5 | 893.1 | 895.8 |
| Biased integration | 1326.7 | 822.1 | 843.2 | 1291.2 | 894.4 | 896.2 |
| a,b from 1finger exp. | **1345.5** | **828.6** | **846.5** | **1308.9** | **899.9** | **913.2** |
| a=1 | **1335.8** | **825.8** | 844.8 | **1301.0** | **901.8** | **905.7** |
| b=0 | **1349.6** | **830.7** | **849.4** | **1301.2** | **899.5** | **902.9** |
| Vector sum. | 1326.7 | 821.8 | 843.2 | 1291.2 | 894.4 | 896.2 |
| No norm. | **1333.8** | **829.2** | **848.5** | **1295.1** | 896.2 | **899.0** |
| Lateral int. | 1327.0 | **825.0** | 844.9 | 1290.5 | 895.2 | 897.2 |
| Lateral inh. | **1334.2** | 820.8 | 842.1 | 1292.1 | 894.5 | **898.6** |

*Note: those with significant difference from the biased integration model are shown in bold fonts. Vector sum. (vector summation model); No norm. (model without normalization); Lateral int. (model with lateral interaction); Lateral inh. (model with lateral inhibition)*

**S11: Weighted averaging of vectors and weighted averaging of angles.** Let's assume that the left-most and reft-hand-side fingers received motion stimuli of angles $\theta_1$ and $\theta_2$, respectively. When we calculate the weighted average of the two motion vectors and determine its angle, the angle $\bar{\theta}_{vec}$ can be given as follows.

$$\bar{\theta}_{vec} = atan\left(\frac{\sum_{i=1}^{2} w_i sin\theta_i}{\sum_{i=1}^{2} w_i cos\theta_i}\right) \tag{S11.1}$$

When the two angles are small this can be simplified as follows.

$$\bar{\theta}_{vec} \approx atan\left(\frac{\sum_{i=1}^{2} w_i \theta_i}{\sum_{i=1}^{2} w_i}\right) \tag{S11.2}$$

Furthermore, since the Maclaurin series of the arc tangent function can be given as follows

$$atan(\,x\,) = x - \frac{x^3}{3} + \frac{x^5}{5} - \frac{x^7}{7} + \cdots \tag{S11.3}$$

equation (S11.2) can be simplified further as follows.

$$\bar{\theta}_{vec} \approx \frac{\sum_{i=1}^{2} w_i \theta_i}{\sum_{i=1}^{2} w_i} = \bar{\theta}_{ang} \tag{S11.4}$$

Note that this is equivalent to the weighted average of the individual angles ($\bar{\theta}_{ang}$).

Finally, we also compared $\bar{\theta}_{vec}$ and $\bar{\theta}_{ang}$, calculated from equations (S11.1) and (S11.4), for all the stimulus pairs with weight ranging from 0 to 1. The maximum difference between the two angles was below 0.15 degrees.
